# MSAP in Tiger Snakes: Island populations are epigenetically more divergent

**DOI:** 10.1101/118836

**Authors:** Carlos Marcelino Rodríguez López, Moumouni Konate, Vicki Thomson

## Abstract

Research on changes in phenotypic plasticity within wild animal populations is centuries old, however far fewer studies have investigated the role that epigenetics play in the development or persistence of natural variation in response to environmental change. Tiger snakes (*Notechis scutatus*) are an ideal study organism to investigate the link between epigenetics and phenotypic responses to environmental change, as they live on a range of offshore islands with different environments and prey types while exhibiting gigantism and dwarfism in body and head size. In this study, we have generated methylation sensitive amplified polymorphism (MSAP) data and found that, in general, Tiger Snakes are more epigenetically differentiated than genetically differentiated. Each island group has a distinctive epigenetic signal, suggesting the Tiger Snakes on each island group have adapted to their specific environment. This is also supported by the strong positive relationship between epigenetic differentiation and isolation age, as well as between epigenetic/genetic signal and both temperature and precipitation. The Tiger Snakes from Kangaroo Island, which has a complex landscape/environment like the mainland rather than the simple landscape/environment of each of the smaller islands, are both genetically and epigenetically more like the mainland. As the MSAP loci are randomly distributed across the genome, we believe a closer examination of the epigenetic modifications near genes involved in growth, development, and lipid metabolism will allow us to investigate the epigenetic basis for the natural variation in head size and body size on the islands.

## Introduction

Evolutionarily adaptive responses in animals take place through a complex set of mechanisms that includes physical traits, behaviour, and environment but also the genes encoding for these characteristics and their epigenetic regulation. It has long been hypothesised that important fitness traits across a species, such as variation in body size and the tolerance of aridity and heat, may not be hard-wired in the DNA, but rather influenced by epigenetic modification to the DNA backbone that allows immediate responses to environmental cues (Feinberg & Irizarry 2011). Thus the phenotype of an organism appears to be influenced by epigenetic markers to some degree, with aberrations in these mechanisms creating variation that can lead both to adaptive and maladaptive traits (Consuegra and Rodriguez Lopez, 2016). DNA methylation is the best, and most easily, characterized epigenetic modification, and one that has been demonstrated to remain stable through the germline (Jablonka & Raz 2009).

Little is known about epigenetic variation in wild populations, how it is generated and how it relates to phenotypic plasticity. Island populations provide a source of natural variation, which can be used to assess naturally occurring epigenetic variation. An ideal species for examining epigenetic variation in animals in the Australian context is the Tiger Snake, as it occurs on multiple offshore islands that have been isolated for varying amounts of time. The isolation of Tiger Snakes on Australia’s offshore islands stem from both sea-level rise after the last glacial maximum (6-14,000 years ago [ya]) and the transport and release of captive populations by humans ~100 ya. In the recently isolated populations, neonate Tiger Snakes can adjust their head size to suit prey size (Aubret & Shine 2009), suggesting an epigenetic regulation of growth genes. Whereas on island populations with large prey items separated for thousands of years, the Tiger Snakes are born, and/or develop into, giant-sized snakes (Shine 1987), suggesting the traits governing body/head gigantism have been genetically assimilated. This well-characterised phenotypic plasticity makes the Tiger Snake an ideal wild species to investigate the link between epigenetic gene regulation and phenotypic plasticity.

In this study, we use methylation sensitive amplified polymorphism (MSAP) to characterise the differences in methylation between Tiger Snake populations and correlate with various population characteristics, such as geographic distance between populations, isolation age, and bioclimatic variables.

## Methods

### DNA extractions

A total of 70 samples were obtained from the Australian Biological Tissue Collection (ABTC) at the South Australian Museum (SAM) or from the Australian National Wildlife Collection (ANWC) at the Commonwealth Scientific and Industrial Research Organisation (CSIRO). Sample geographic origin was determined from the specimen voucher information. In brief, a total of 38 samples were originally collected from island populations, including Kangaroo Island, Chappell Island, Nuyts Archipelago (Goat, Franklin and St Peter Islands) and Sir Joseph Banks Islands (Roxby Island, Reevesby Island and Hareby Islands) while 32 samples where from continental populations (for detailed information see Supplementary Table 1). DNA was extracted from the tissue samples using either a ‘salting-out’ method (Nicholls *et al.* 2000) or a DNeasy Blood and Tissue kit (Qiagen).

### Amplified Fragment Length Polymorphism (AFLP) and Methylation Sensitive Amplification Polymorphism (MSAP) Profiling

AFLP and MSAP methods were adapted from (Rodríguez Lopez *et al.* 2012). Briefly, 5.5 μl of the normalized gDNA were used for parallel DNA restriction and ligation performed using a combination of restriction enzyme EcoRI and *Mse*I for AFLPs and *EcoR*I and one of two isoschizomers *Msp*I or *Hpa*II for MSAPs. DNA was added to 5.5 μl the Restriction/Ligation master mix containing: 1.1 μl T4 Ligase buffer (×10), 1.1 μl Nacl (0.5 M), 0.55 μl BSA (1 mg/ml), 1U *Mse*I (New England Biolabs) or *Hpa*II (New England Biolabs) or MspI (New England Biolabs), 5U *EcoR*I (New England Biolabs), 20U T4 DNA Ligase (New England Biolabs), 1 μl *EcoR*I adapter (5 μM), 1 μl *Hpa*II*/Msp*I adapter (50 μM) or 1 μl MseI adapter (50 μM) (See Table 1 for full sequences), and water to a final reaction volume of 11 μl. Restriction/Ligation was performed using a T100^TM^ Thermo cycler (Bio-Rad Laboratories, Hercules, CA) with the following protocol: 2 h at 37°C followed by 10 min at 65°C.

**Table 1.**
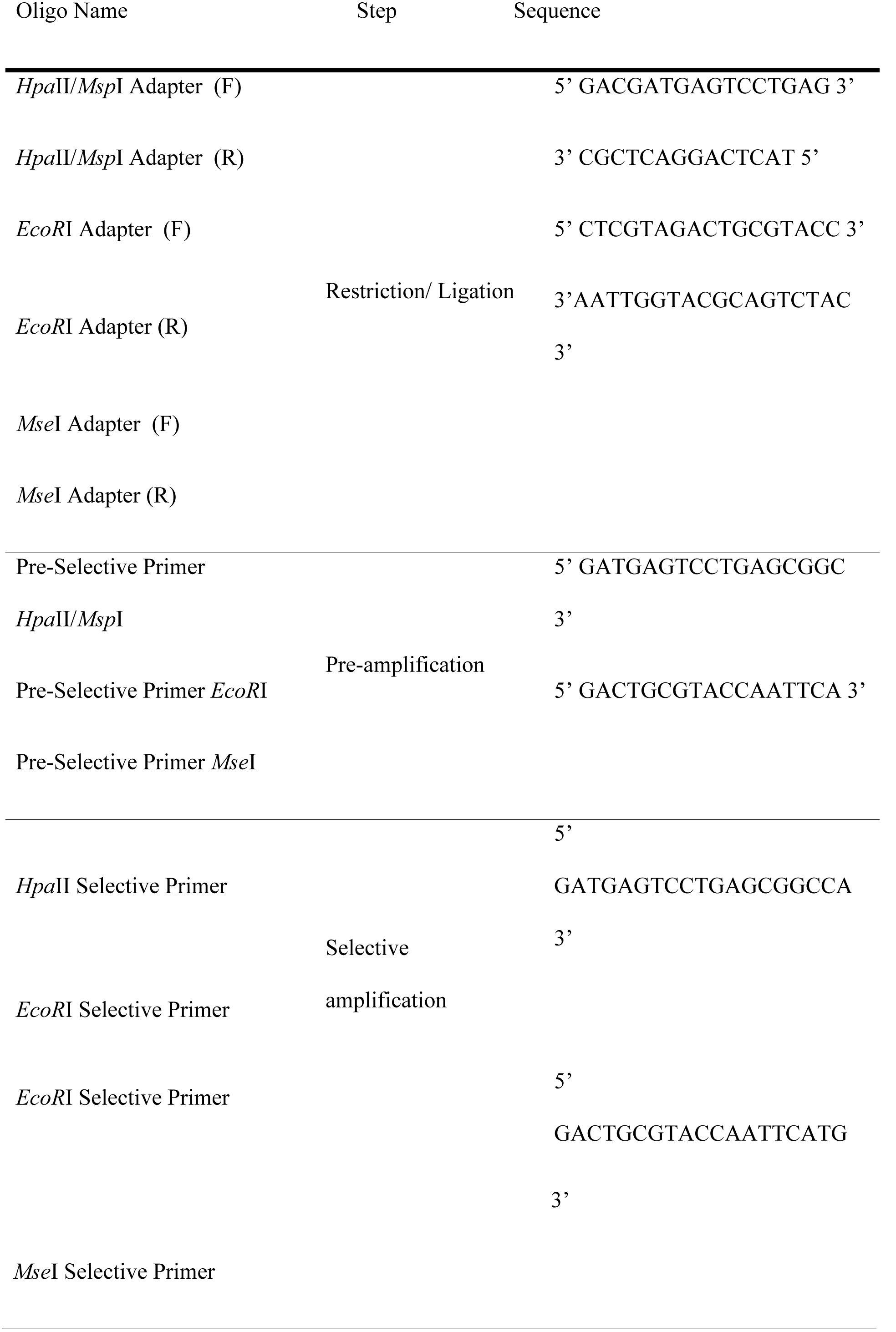
Oligonucleotide Sequences used for MSAP

For the pre-amplification step, 1 μl of each Restriction/Ligation reaction product was added to 11.5 μl of pre-amplification master mix containing: 6.25 μl 2x Biomix (Bioline), 0.25 μl *Hpa*II*/Msp*I pre-amplification primer (10 μM) or 0.25 μl MseI pre-amplification primer (10 μM), 0.5 μl *EcoR*I pre-amplification primer (10 μM) (Table 1), 0.1 μl BSA (1 mg/ml) and 4.85 μl Water. PCR amplification was performed using a T100^TM^ Thermo cycler (Bio-Rad Laboratories, Hercules, CA) with the following protocol: 2 min at 72°C, followed by 30 cycles of 30 sec at 94°C, 30 sec at 56°C, 2 min at 72°C with a final step of 10 min at 72°C.

A second round of selective amplification was carried out by adding 1 μl of the Pre-amplification PCR product to 11.5 μl of the Selective amplification Master mix containing: 6.25 μl 2x Biomix (Bioline), 0.25 μl 5’-FAM labelled *Hpa*II/*Msp*I Selective primer (10 μΜ) or 0.25 μl 5’-FAM labelled *Mse*I Selective primer (10 μΜ), 0.5 μl *EcoR*I Selective primer (10 μΜ) (Table 1), 0.1 μl BSA (1 mg/ml) and 4.85 μl nanopure water. PCR amplification was performed using a T100™ Thermo cycler (Bio-Rad Laboratories, Hercules, CA) with the following protocol: 2 min at 94°C, followed by 13 cycles of 30 sec at 94°C, 30 sec at 65°C (reduce by 0.7°C per cycle), 2 min at 72°C, followed by 24 cycles of 30 sec at 94°C, 30 sec at 56°C, 2 min at 72°C with a final step of 10 min at 72°C. Selective amplification products were sent for capillary electrophoresis separation to the Australian Genome Research Facility Ltd (AGRF), Adelaide, South Australia.

### Statistical analysis

MSAP profiles were visualized using GeneMapper Software v4 (Applied Biosystems, Foster City, CA). Binary matrices containing presence (1) absence (0) allelic information were generated from the capillary separation results obtained from samples restricted using enzyme combinations *Eco*RI/*Mse*I (AFLPs) and *Eco*RI/*Msp*I and *Eco*RI*/Hpa*II (MSAPs). In this case, fragment selection was limited to allelic sizes between 85 and 550 bp to reduce the potential impact of size homoplasy (Caballero *et al.* 2008). In all cases, different levels of hierarchy were generated to group the samples. Samples were first grouped into continental or island populations. Then, samples from island populations were divided according to the island they were collected from. Finally, samples from island populations were separated by island age.

The numbers of the various fragments attributed to non-methylated (+/+), CHG methylated (+/-), CG methylated (-/+), and uninformative (-/-) were calculated for each population based on the MSAP profiles. Shannon Diversity indices were used to estimate the within-population (at the island group level) epigenetic diversity (H_pop_) of the CG and CHG methylation patterns and the Kruskal-Wallis H test was used to estimate the significance of the differences in Shannon diversity index among these populations.

For the analysis of the AFLP and MSAP data, GenAlex v6.5 software (Peakall & Smouse 2012) was used to infer pairwise genetic (PhiPT calculated from AFLP profiles) and epigenetic (ePhiPT calculated from MSAP profiles) distances between the different hierarchical groups described above. Analysis of Molecular Variance (AMOVA) was then performed using the same software to test the significance of the estimated distances between groups using 9999 random permutations. Principal Coordinates Analysis (PCoA) was used to visualize the patterns of epigenetic variation associated to geographical origin and island age. The PCoA were performed in the R package ‘stats’ using the prcomp command.

Mantel test analyses were then used to estimate the correlation between: 1) the calculated pairwise molecular distances (i.e. PhiPT and ePhiPT) and the pairwise geographic distance in km (GGD) among populations and 2) the calculated pairwise molecular distances (i.e. PhiPT and ePhiPT) and the pairwise age differences among populations as described by Rois *et al.* (2013). Mantel test significance was assigned as the probability of the correlation between compared matrices using random data being higher than the correlation between matrices of collected data (i.e. P(rxy-rand >= rxy-data)) estimated over 9,999 random permutations tests, as implemented in Genalex v6.5.

### Climate

The 19 bioclim layers were downloaded from the worldclim website (worldclim.org/bioclim), converted from tiff to ascii files and trimmed to focus on Australia using the R computing package Raster. The values of the 19 bioclim layers for each of the South Australia sample locations used in this study were extracted from the climate data and input into a principal component analysis (PCA) to examine how each population varies in overall climate. Mantel test analyses (R package ade4) were also used to estimate the correlation between differences in epigenetic/genetic principal components (i.e. PC1 of the MSAP PCoA) and each of the 19 bioclimatic variables for the sampling locations within South Australia (where most of the samples were collected from) and converted to a distance matrix. The relationships between these raw datasets were also plotted with regression lines and Pearson correlation coefficients were calculated to estimate the strength of these relationships using the R package ‘stats’.

## Results

67 Tiger Snake samples were successfully genotyped, with 38 from island populations and 29 from mainland Australia. As there were not enough successfully genotyped samples from some locations to generate population level data, the majority of analyses were performed on samples from South Australia: the Nuyts Archipelago (n=12), Sir Joseph Banks Island group (n=11), Kangaroo Island (n=7) and the South Australian mainland (n=17; see Figures 1 & 2 for sampling locations).

**Figure 1.**
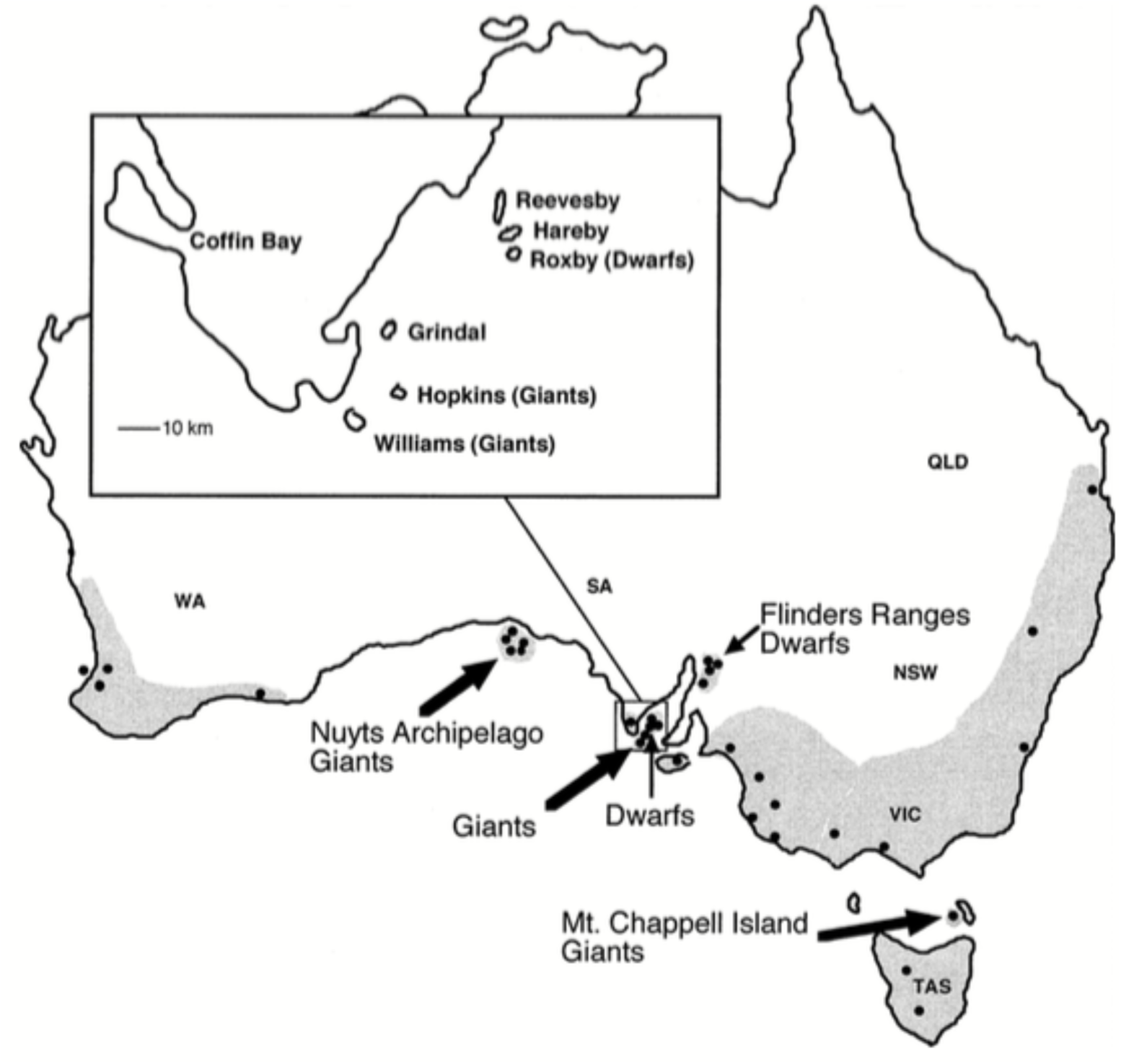
Map of Australia showing Tiger Snake populations sampled in this study (redrawn from Keogh *et al.* 2005).

**Figure 2.**
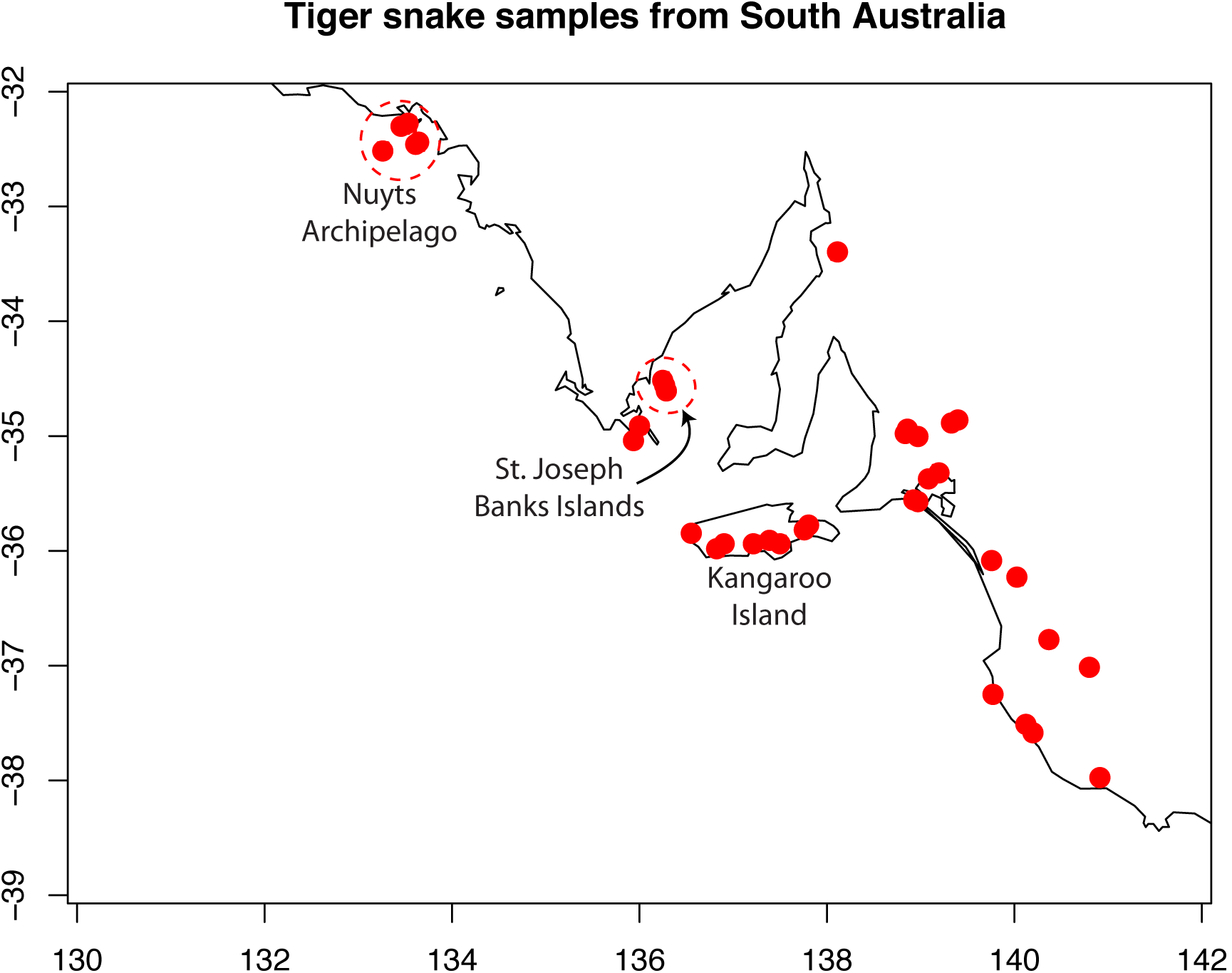
Map of South Australia showing sampling locations and island groups.

The numbers of the various fragments attributed to non-methylated (+/+), CHG methylated (+/-), CG methylated (-/+), and uninformative (-/-) were calculated for the mainland populations and island populations separately based on the MSAP profiles. Within South Australia (SA), the total 5’-CCGG-methylation level (both CG and CHG together) ranged from 22% (of fragments) in mainland SA Tiger Snakes to 30% on Roxby Island (Table 2 & Figure 3). The amongst SA population difference in overall genome-wide methylation levels (both CG and CHG methylation together) was not significant (ANOVA p = 0.6270), nor was the difference in CHG methylation between SA populations (ANOVA p = 0.1637); however the difference in CG methylation between each of the SA populations was significant at the 5% level (ANOVA p = 0.0341; range from 10% of fragments on Goat Island to 17% on Roxby Island; Table 2). Shannon diversity indices were calculated for each methylation type per SA population group (Table 3) and Kruskal-Wallis rank sum tests were used to test for differences between Tiger Snakes on the SA mainland and each of the island groups, which were significantly different at the 10% level for CG methylation (chi-squared = 2.9503, p = 0.08586), and for CHG methyation (chi-squared = 3.3103, p = 0.06884). The coefficient of epigenetic differentiation was also higher in the SA mainland populations compared to the SA island populations (Table 4.)

**Table 2.**
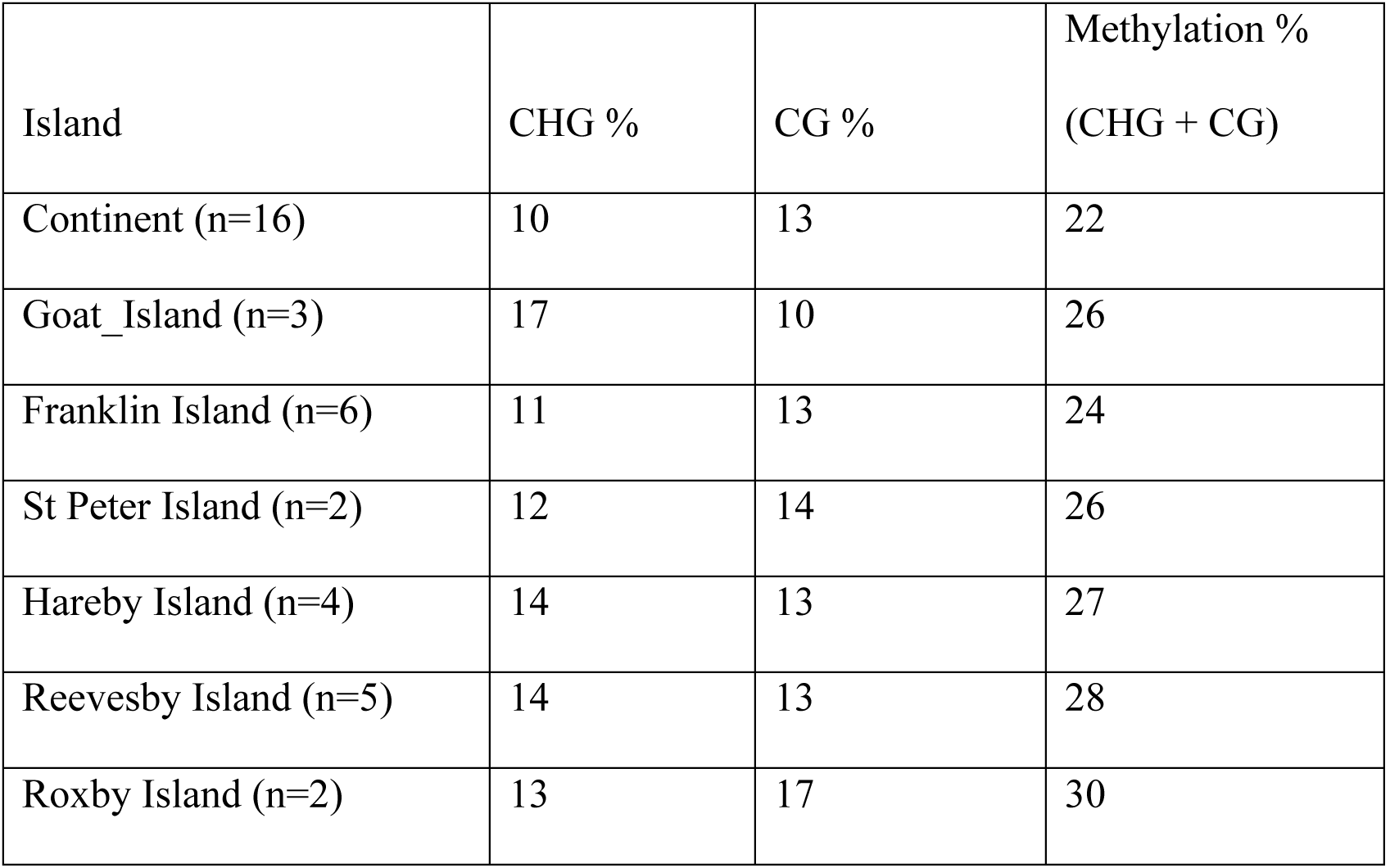
Each type of methylation as a percentage of fragments genotyped for the SA populations.

**Figure 3.**
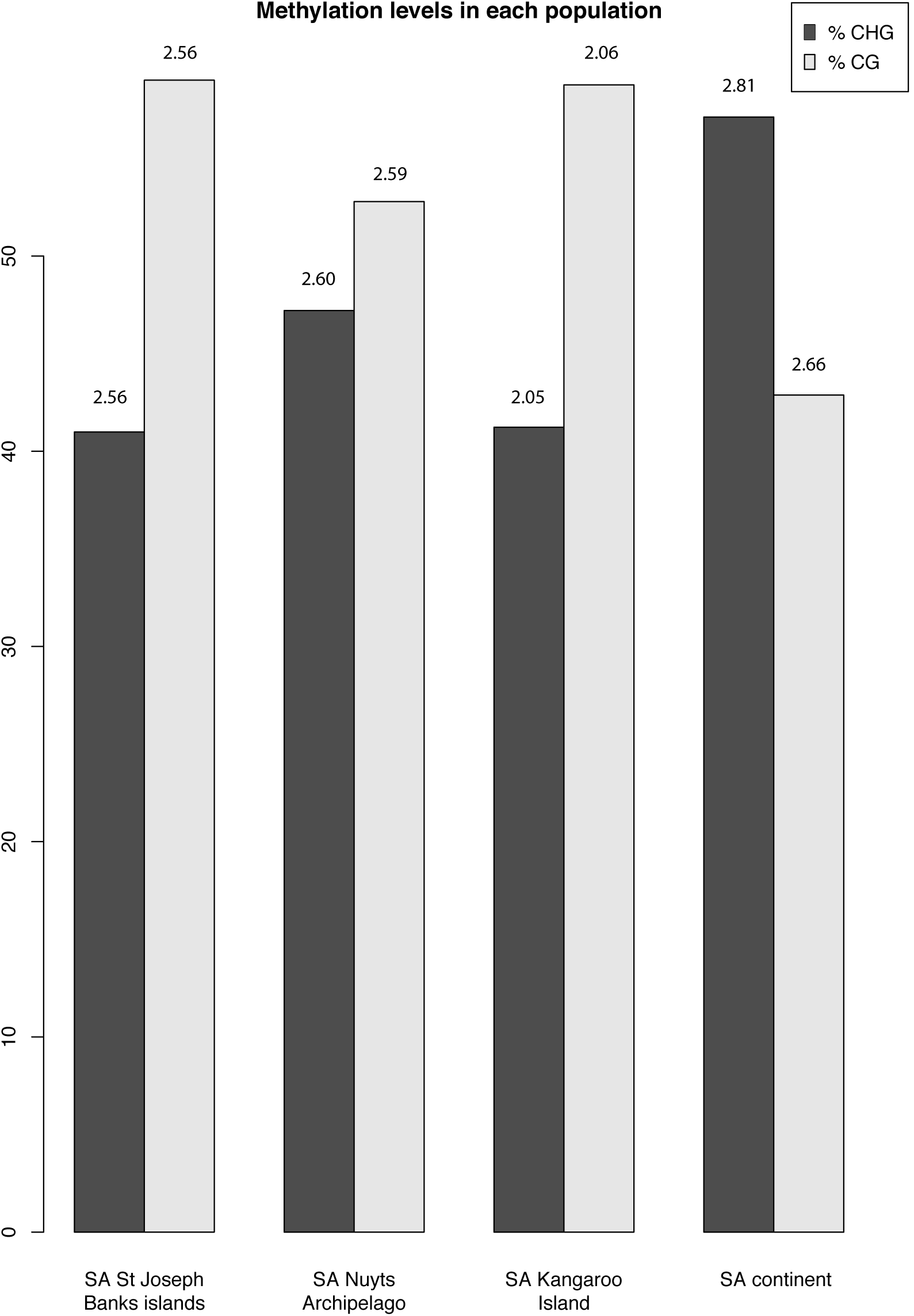
Histogram of CG and CHG methylation levels as a percentage of the total methylated fragments in the South Australia mainland vs. island Tiger Snakes. Values on top of the histogram bars are the Shannon Diversity Indices for within-population epigenetic diversity (H_pop_).

**Table 3.** Shannon Diversity indices used to estimate the within-population epigenetic diversity (H_pop_).

|  | SA<br>mainland | Nuyts<br>Archipelago | Sir Joseph<br>Islands | Kangaroo<br>Island | Species-<br>level |
| --- | --- | --- | --- | --- | --- |
| Shannon<br>diversity index<br>(CHG) | 2.8188 | 2.6003 | 2.5536 | 2.0440 | 3.9222 |
| Shannon<br>diversity index<br>(CG) | 2.6625 | 2.5764 | 2.5584 | 2.0425 | 3.8699 |

**Table 4.** Measures of genetic distance for the AFLP data between the South Australian populations, with values below the diagonal pairwise representing Nei’s genetic distance and above the diagonal are pairwise PhiPT (asterisk shows those values significant at the 5% level after 9999 permutations).

|  | KI<br>(n=7) | Mainland<br>(n=17) | Nuyts<br>(n=12) | Sir Joseph<br>(n=11) |
| --- | --- | --- | --- | --- |
| KI | - | 0.024 | 0.124 * | 0.080 * |
| Mainland | 0.034 | - | 0.166 * | 0.090 * |
| Nuyts | 0.061 | 0.072 | - | 0.041 |
| Sir Joseph | 0.050 | 0.047 | 0.031 | - |

### Genetics

The genetic PCoA shows that some Kangaroo Island Tiger Snake samples are genetically closer to mainland samples, while others are genetically closer to island populations (Figures 4 & 5). Within SA the populations that are genetically closest to each other are the Sir Joseph Banks Island group and Nuyts Archipelago populations (Nei genetic distance = 0.031; PhiPT = 0.041; Table 5) and they appear intertwined on the PCoA (Figure 5). Using Nei’s genetic distance, the Kangaroo Island and mainland Tiger Snakes are also genetically quite close (Nei genetic distance = 0.034; PhiPT = 0.024), with the most genetically distant populations the mainland-Nuyts Archipelago populations (Nei genetic distance = 0.072; PhiPT = 0.166).

**Figure 4.**
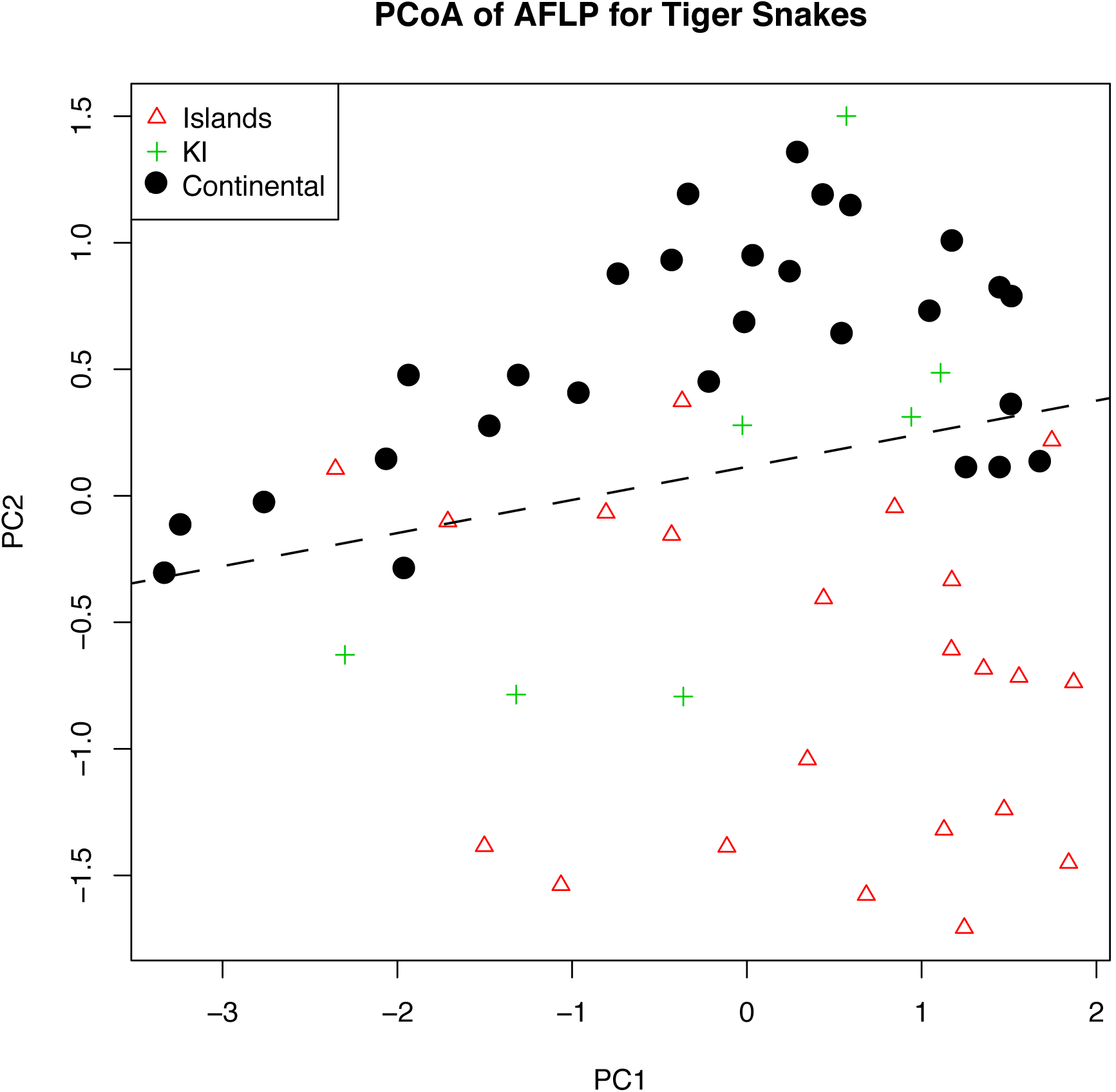
Principal co-ordinate analysis of genetic data (AFLP) for all Tiger Snake samples, showing the major differentiation in PC2 of mainland (black circles) vs. island (red triangles) populations. Some Kangaroo island samples (green crosses) fall with the island populations but others fall with the mainland samples.

**Figure 5.**
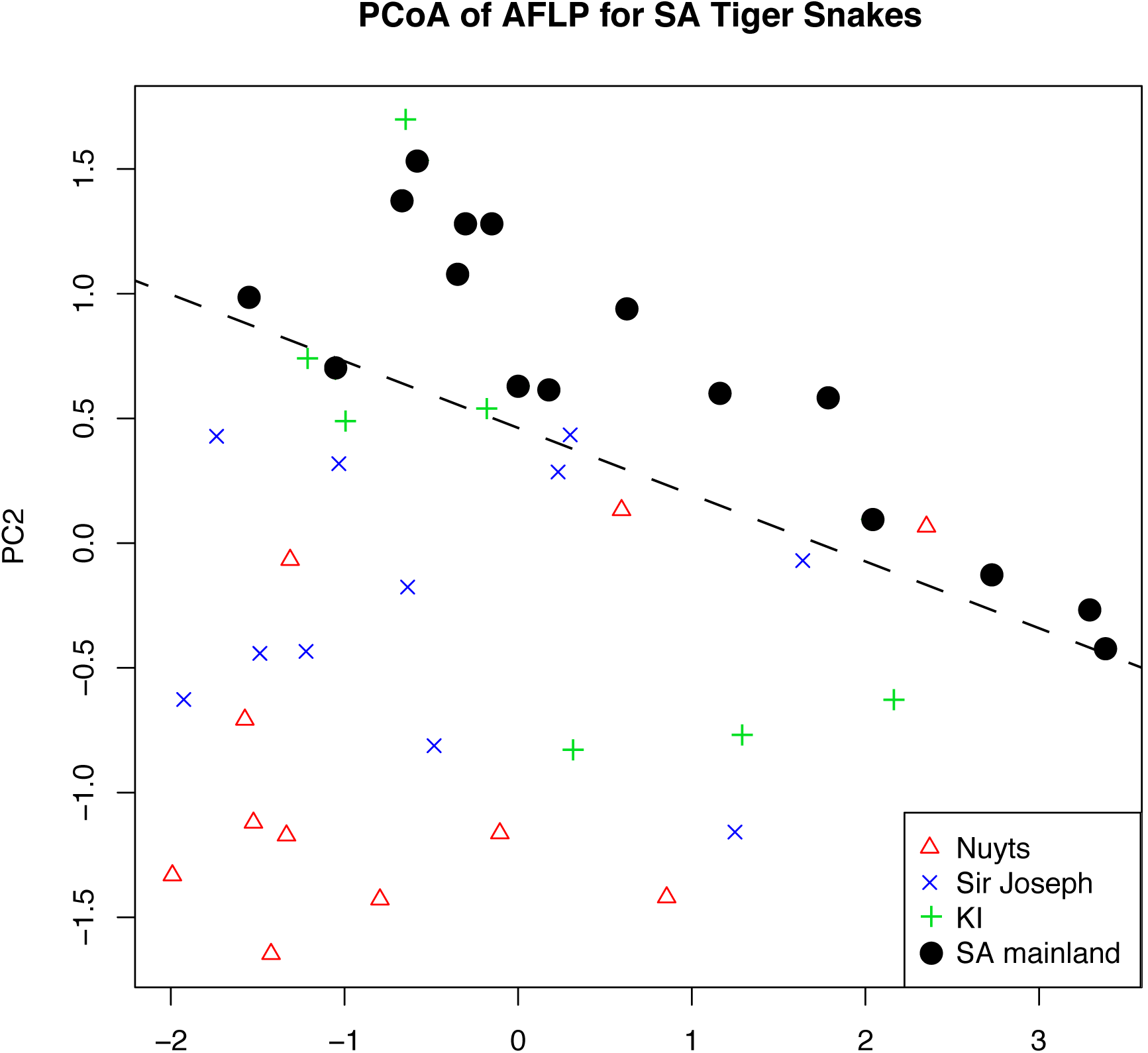
Principal co-ordinate analysis of genetic data (AFLP) for South Australian Tiger Snake samples only (n=59), showing the major differentiation in PC2 of mainland (green plus sign) vs. Sir Joseph Banks island group (blue cross) vs. Nuyts Archipelago (red triangle) populations. Again, some Kangaroo island samples (black circle) fall with the other two island group populations but others fall with the mainland samples.

**Table 5.** The coefficient of epigenetic differentiation was computed as Gst = (Hsp – Hpop)/Hsp where Hsp is the within-species epigenetic diversity.

| Gst | Mainland | Island |
| --- | --- | --- |
| CHG | 0.281311102 | 0.100645628 |
| CG | 0.311996072 | 0.091400959 |

### Epigenetics

The populations that are epigenetically closest to each other are the mainland/KI populations (using both Nei’s and PhiPT epigenetic distance for the HpaII dataset) and the Sir Joseph Banks/Nuyts Archipelago populations (Nei’s and PhiPT epigenetic distance for the MspI dataset; Tables 6 & 7). The populations that are epigenetically most distant are the Sir Joseph Banks islands vs. mainland *(Msp*I PhiPT epigenetic distance) and the Sir Joseph Banks islands vs. all the other populations (HpaII PhiPT epigenetic distance). The Sir Joseph Banks Island group are comprised of Roxby Island, Reevesby Island and Hareby Islands, which each also had the three highest overall percentages of fragments that were methylated (Table 3). The epigenetic PCoA of all Tiger Snake samples, again shows how some Kangaroo Island samples resemble those on the mainland, but others resemble those on the smaller nearby islands (Figure 6).

**Table 6.**
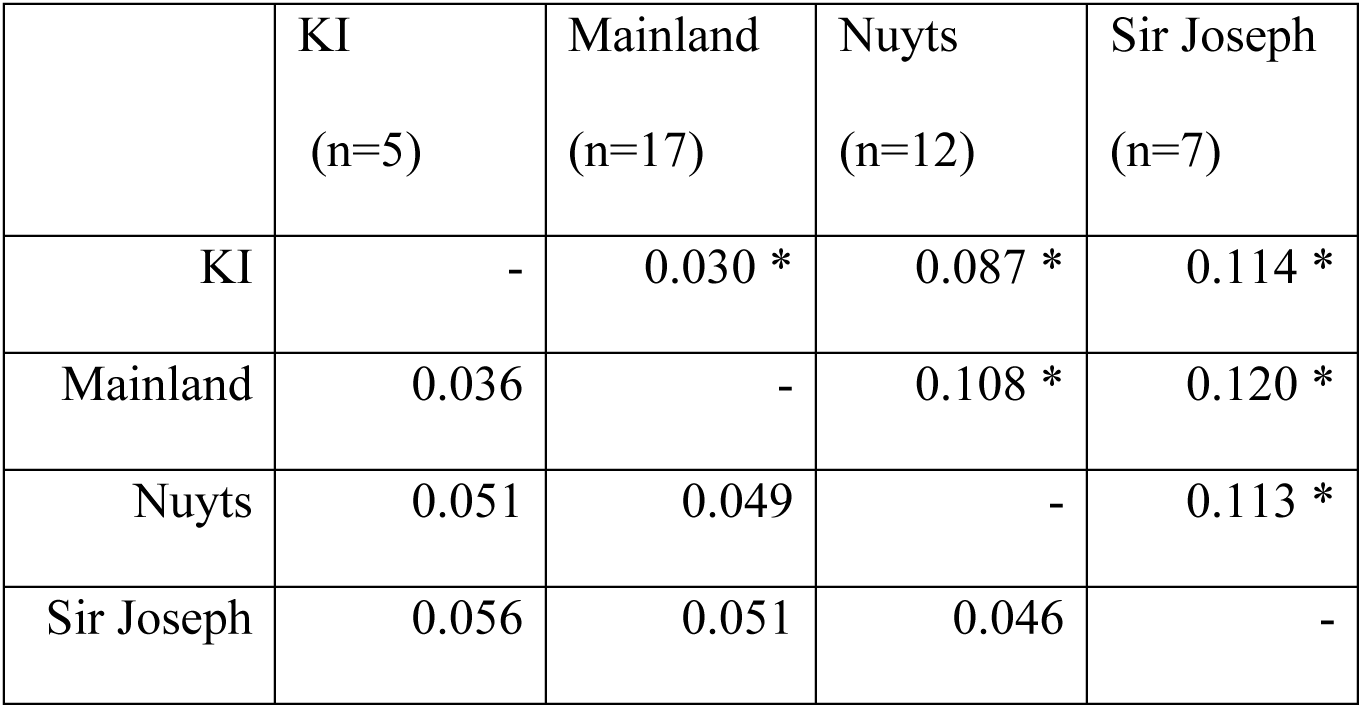
Measures of epigenetic distance for the *Hpa*II data between the South Australian populations, with values below the diagonal pairwise representing Nei’s genetic distance and above the diagonal are pairwise PhiPT (asterisk shows those values significant at the 5% level after 9999 permutations).

**Table 7.** Measures of epigenetic distance for the *Msp*I data between the populations, with values below the diagonal pairwise representing Nei’s genetic distance and above the diagonal are pairwise PhiPT (asterisk shows those values significant at the 5% level).

|  | KI<br>(n=7) | Mainland<br>(n=17) | Nuyts<br>(n=12) | Sir Joseph<br>(n=11) |
| --- | --- | --- | --- | --- |
| KI | - | 0.097 * | 0.090 * | 0.083 * |
| Mainland | 0.055 | - | 0.067 * | 0.106 * |
| Nuyts | 0.050 | 0.037 | - | 0.061 * |
| Sir Joseph | 0.047 | 0.050 | 0.035 | - |

**Figure 6.**
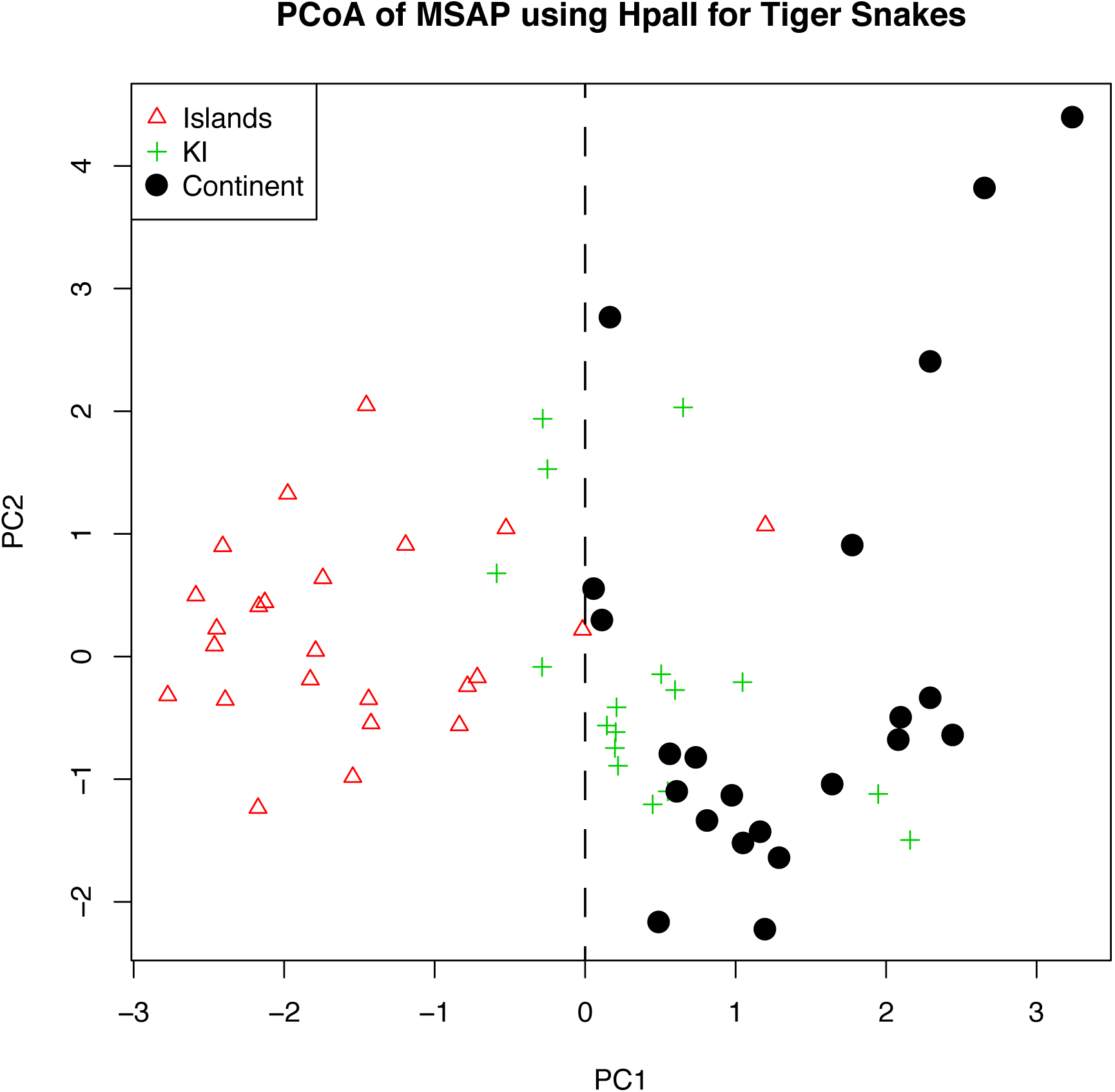
Principal co-ordinate analysis of the epigenetic markers *(Hpa*II) for all the Tiger Snake samples (n=70), showing the first and second principal components.

When only South Australian (SA) Tiger Snakes were examined, the SA mainland samples overlapped to a large degree with those on Kangaroo Island and were quite distinct from those on the Nuyts Archipelago and Sir Joseph Banks Island group in PC1 (Figure 7). Values in PC2, on the other hand, separated islands from the Nuyts Archipelago from those in the Sir Joseph Banks Island group, with the mainland and Kangaroo Island populations in between (Figure 8).

**Figure 7.**
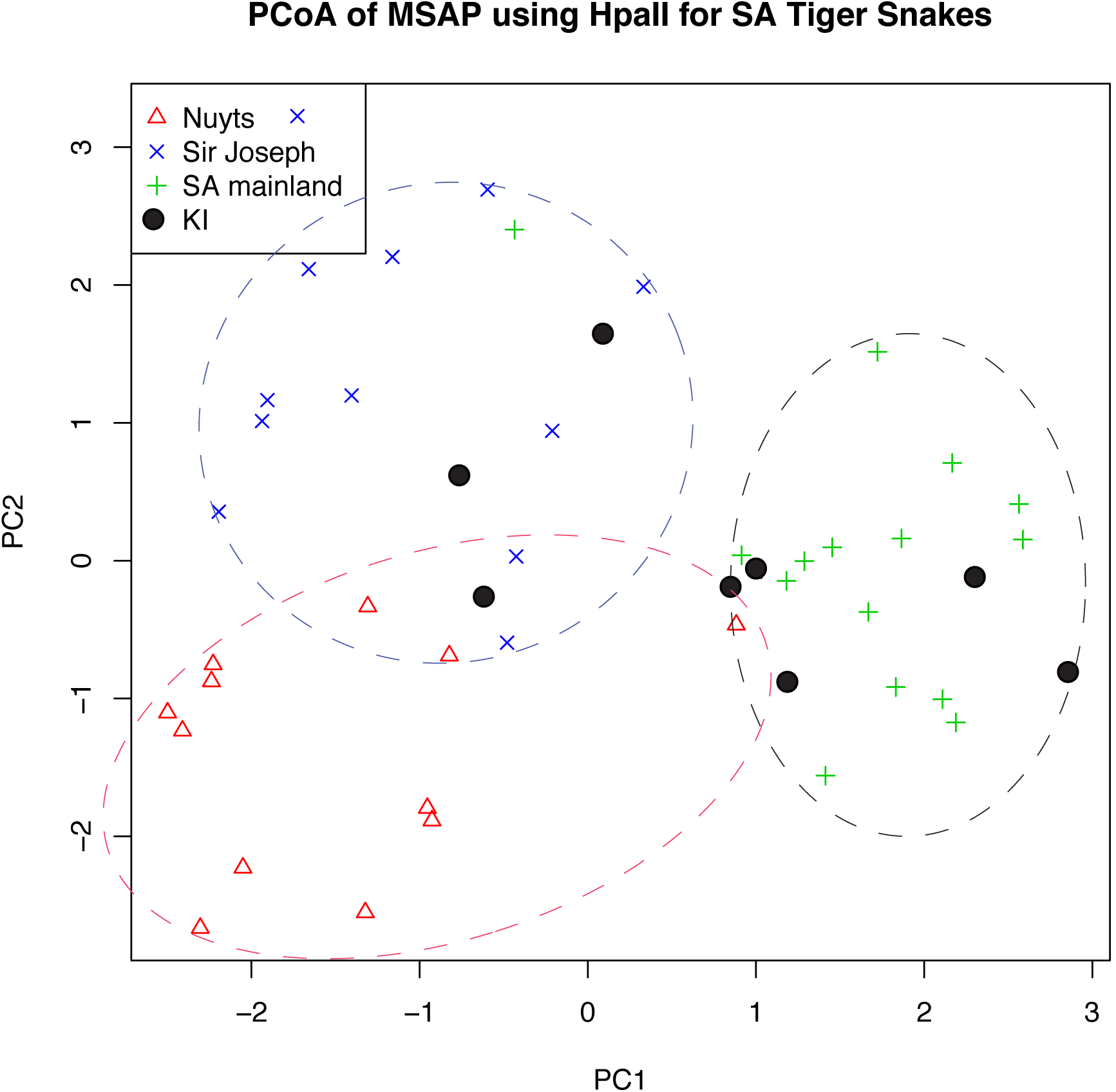
Principal co-ordinate analysis (PCoA) of the MSAP markers for the Tiger Snake samples from South Australia (n=51), showing the first and second principal co-ordinates. PC1 values from this PCoA were used in the mantel and correlation tests to assess the relationships between the epigenetic/genetic signal and bioclimatic variables.

**Figure 8.**
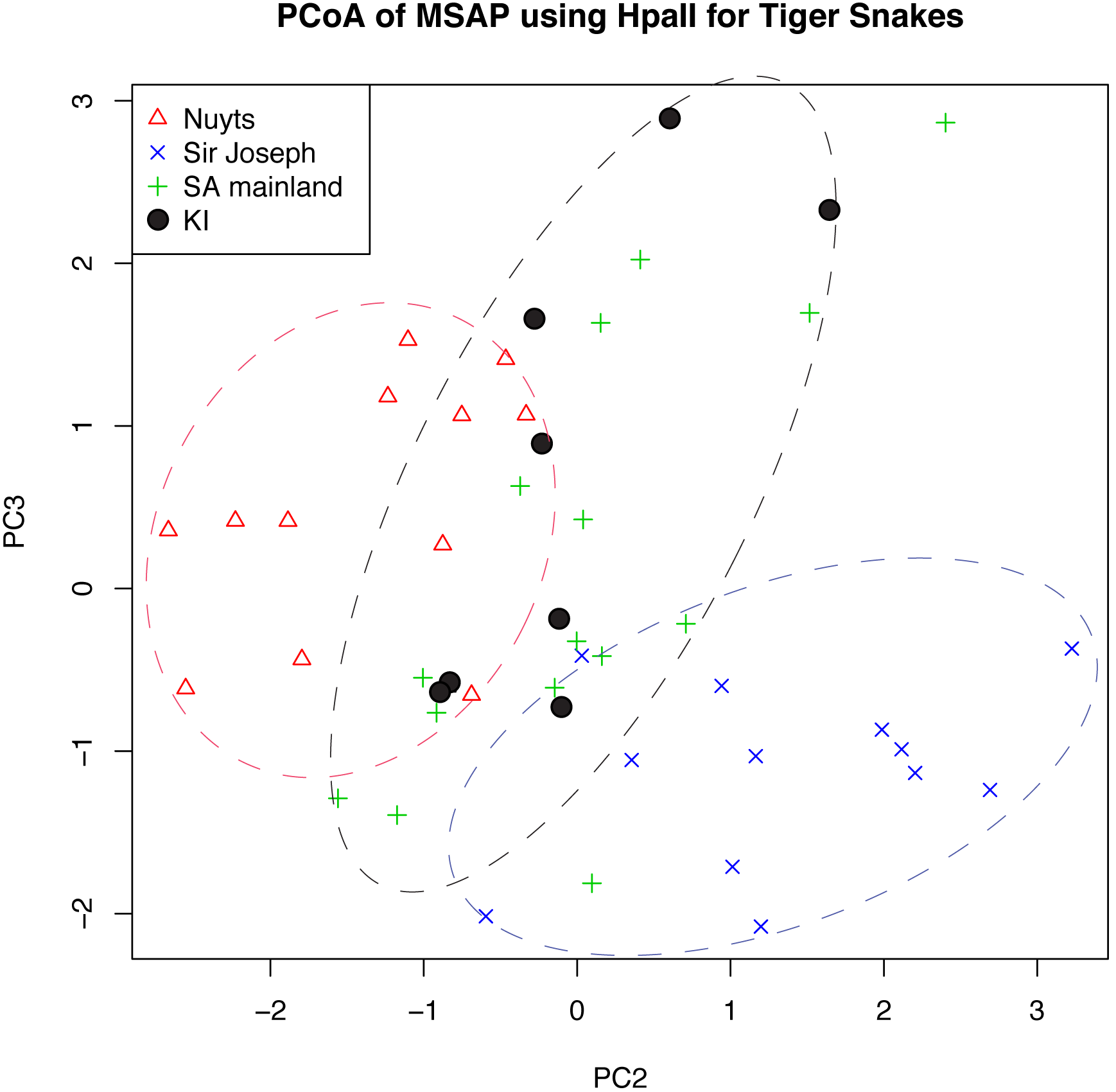
Principal co-ordinate analysis of the MSAP markers for the Tiger Snake samples from South Australia (n=51), showing the second and third principal co-ordinates.

There is also strong correlation (R^2^ = 0.4797 in Figure 9) between genetic distance and geographic distance (traditionally termed isolation-by-distance), however the correlation between epigenetic distance and geographic distance is stronger (R^2^ = 0.5468 in Figure 10). An even stronger correlation also occurs between epigenetic distance and isolation age (R^2^ = 0.5673 in Figure 11 and R^2^ = 0.6922 in Figure 12).

**Figure 9.**
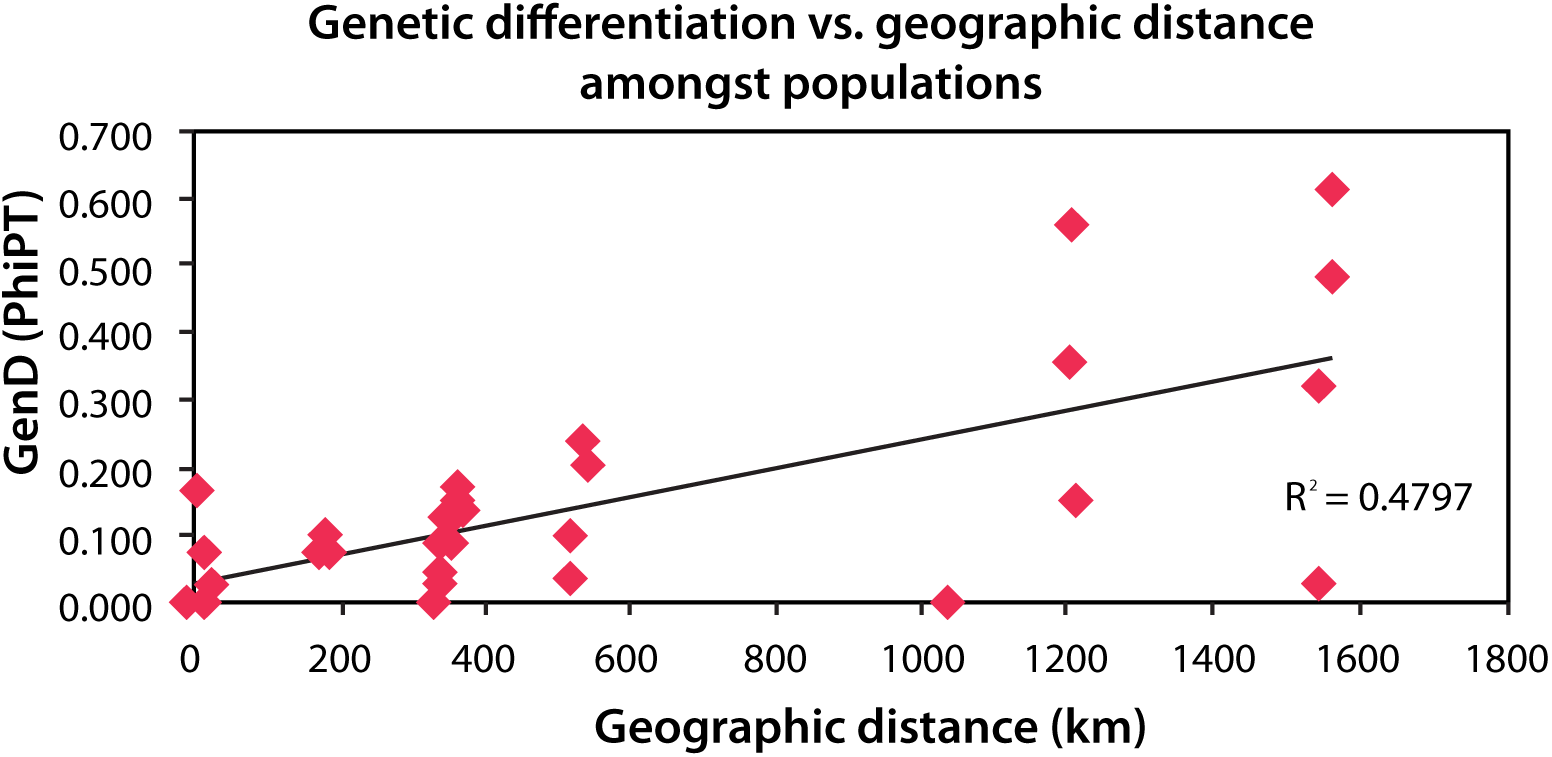
Mantel test of genetic differentiation against geographic distance amongst all the populations (mantel p-value = 0.010).

**Figure 10.**
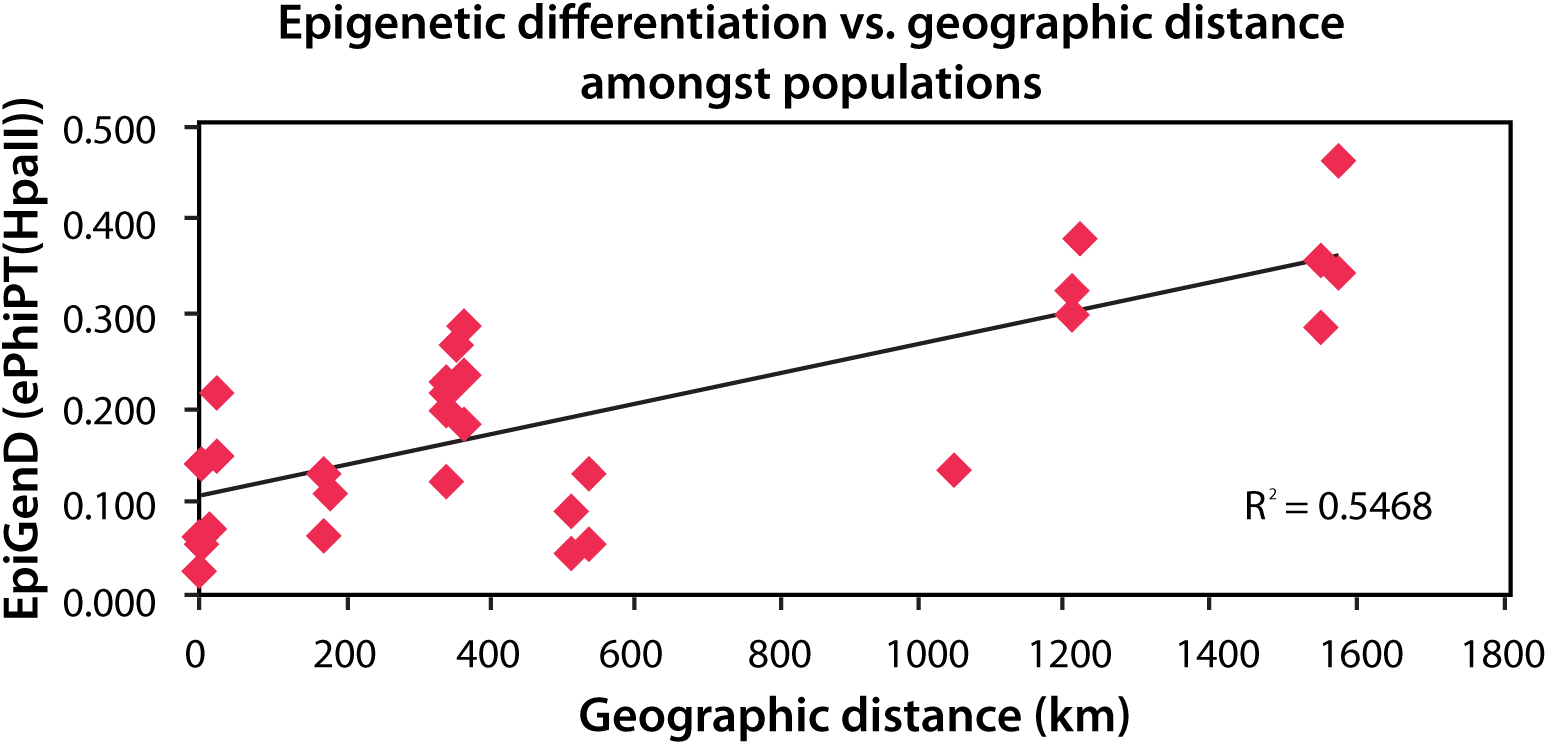
Mantel test of epigenetic differentiation against geographic distance amongst all the populations (mantel p-value = 0.010).

**Figure 11.**
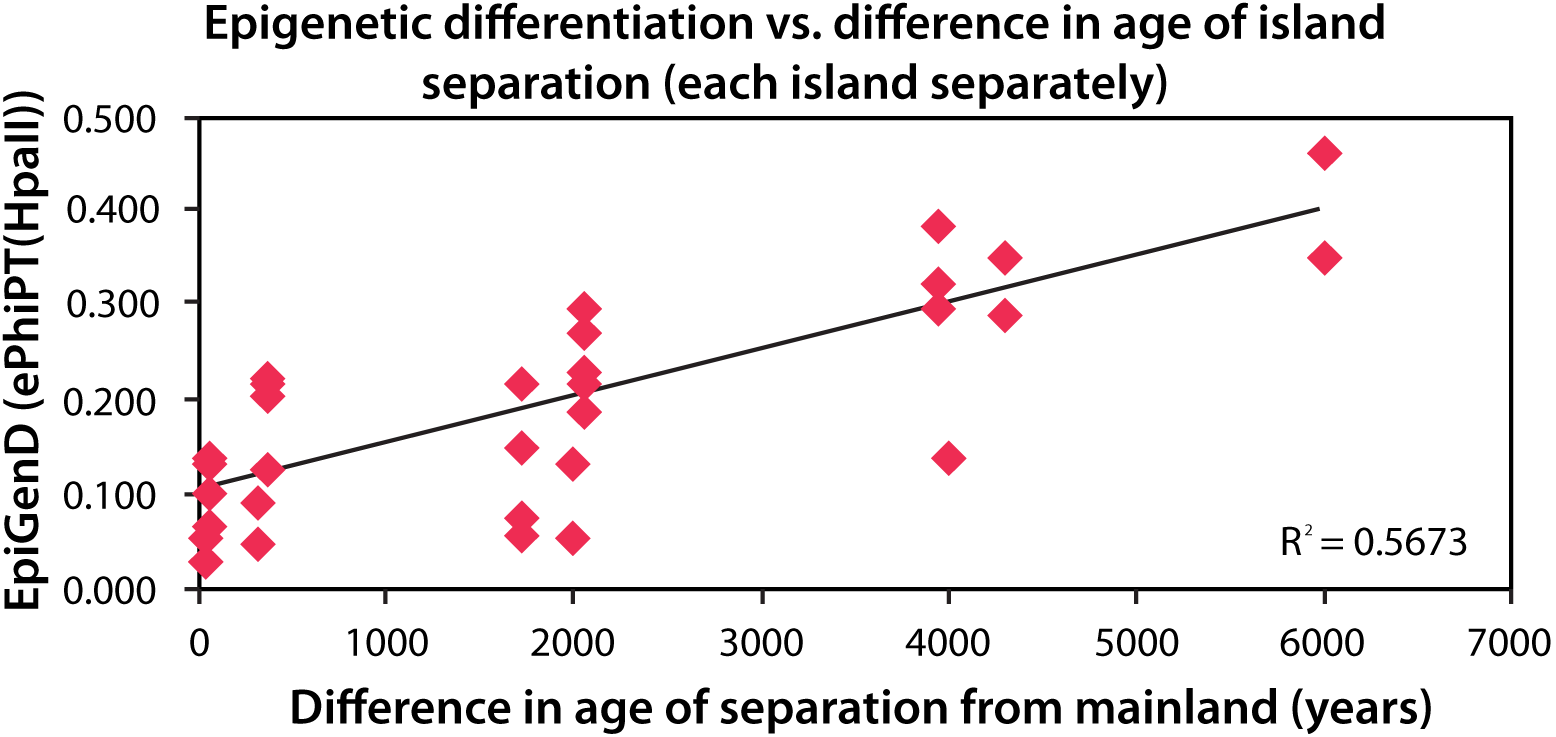
Mantel test of epigenetic differentiation against time since isolation between all the populations (mantel p-value < 0.001).

**Figure 12.**
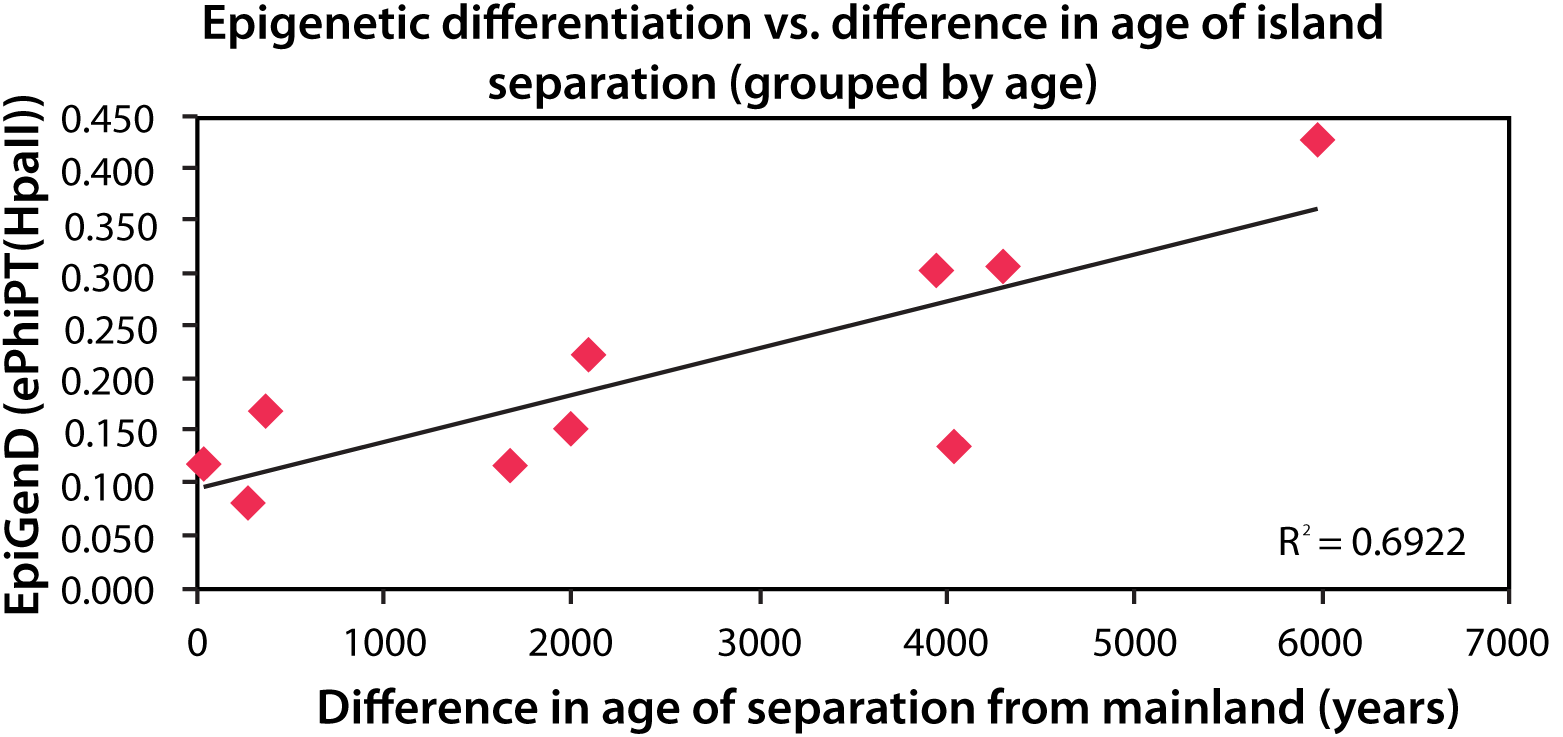
Mantel test of epigenetic differentiation against time since isolation between all the populations grouped by age (mantel p-value = 0.015)

### Environmental data

There are some clear environmental differences between some of the Tiger Snake populations (Figures 13 & 14). The two main bioclimatic variables driving the PCA relationships between the populations are BIO4 (temperature seasonality) and BIO12 (annual precipitation). When we look at how these bioclimatic variables are related to epigenetic distance, we find that similar to the strong correlation between epigenetic distance and isolation age, there are also even stronger correlations between epigenetic distance and bioclimatic variables for the South Australian (SA) samples. The SA mainland samples generally experience higher annual mean temperatures (~15.7—17.3°C) compared to Tiger Snakes on the Nuyts Archipelago and Sir Joseph Banks islands (~13.5—16.3°C), which appears to be strongly related (r = 0.652) to their genetic/epigenetic profiles *(Hpa*II PCoA1 values; Figure 15 & 16). The SA mainland samples generally experience lower precipitation levels in the driest month (< ~12mm) compared to Tiger Snakes on the Nuyts Archipelago and Sir Joseph Banks islands (> ~12mm), which again appears to be strongly related (r = - 0.711) to their genetic/epigenetic profiles *(Hpa*II PCoA1 values; Figures 17 & 18). Other bioclimatic variables related to mean temperature and precipitation levels are also strongly associated with PC1 in the *Hpa*II PCoA, however Bio1 and Bio14 have the strongest correlation (Table 8).

**Figure 13.**
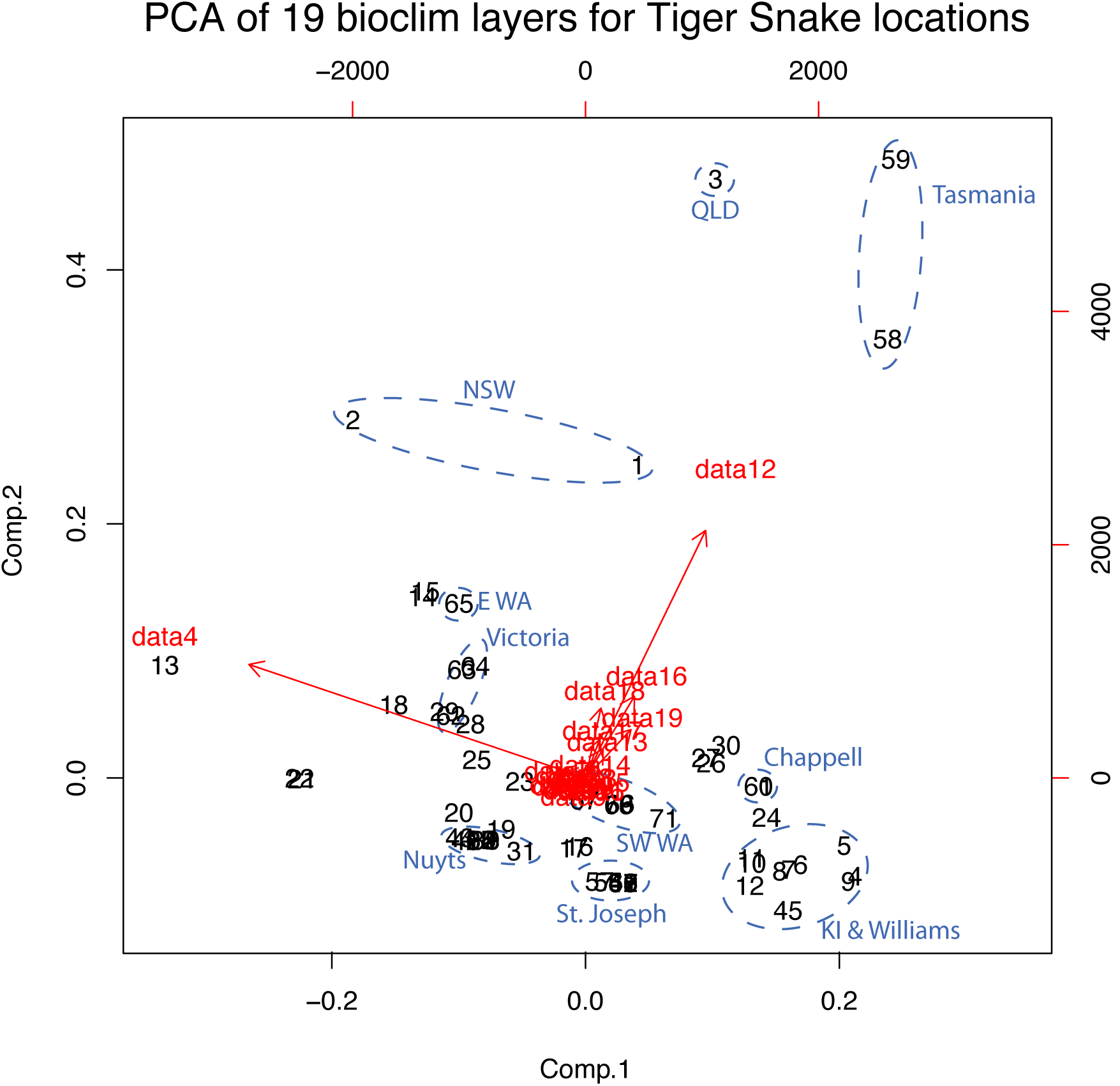
Principal component analysis (PCA) of 19 bioclimatic layer values for the all Tiger Snake sampling locations. The climate layer responsible for most of the variation in principal component 1 is BIO4, which is temperature seasonality, and the climate layer responsible for most of the variation in principal component 2 is BIO12, which is annual precipitation.

**Figure 14.**
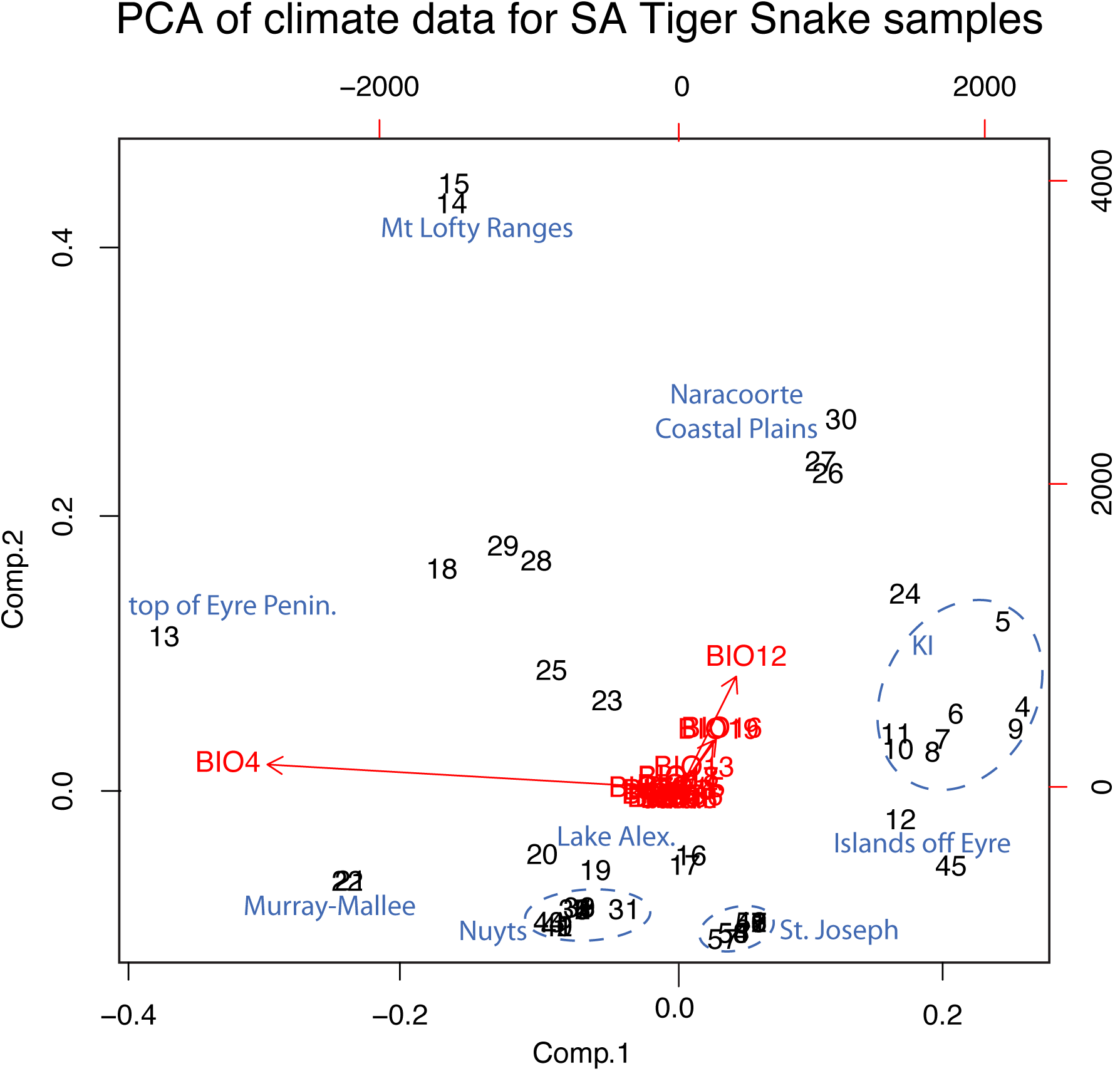
Principal component analysis (PCA) of 19 bioclimatic layer values for the South Australia Tiger Snake sampling locations. The climate layer responsible for most of the variation in principal component 1 is BIO4, which is temperature seasonality, and the climate layer responsible for most of the variation in principal component 2 is BIO12, which is annual precipitation.

**Figure 15.**
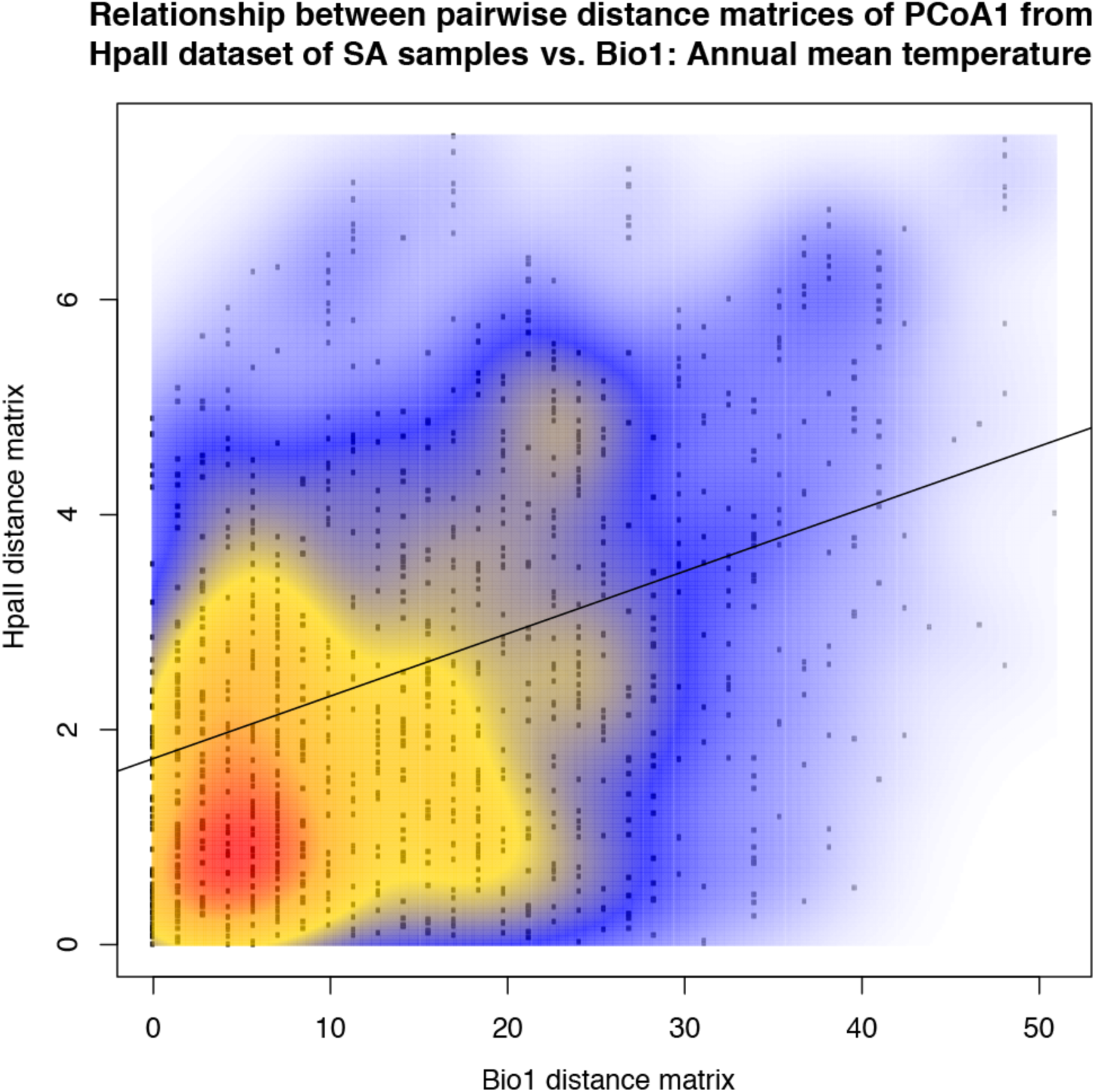
Mantel test showing correlation between pairwise distance matrix of *Hpa*II PCoA1 values and pairwise distance matrix of Bio1 variables (Annual mean temperature) for Tiger Snakes from South Australian locations. The correlation is significantly different from random, with a p-value of 0.001 (Bonferroni corrected p-value of 5% significance level is 0.0026).

**Figure 16.**
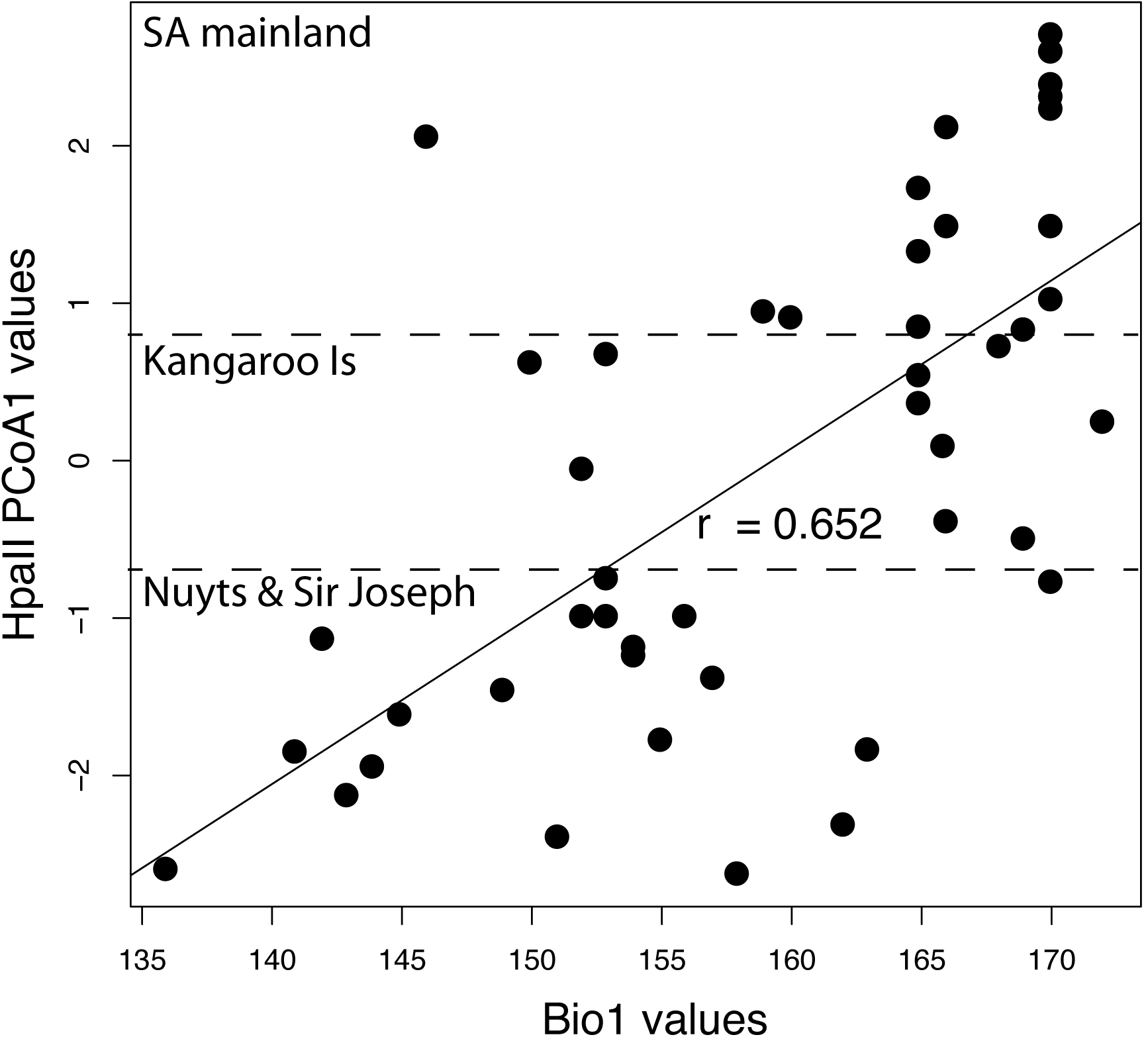
Scatterplot of raw *HpaII* PCoA1 values vs. Bio1 (annual mean temperature) values for Tiger Snakes from South Australian (SA) locations. The horizontal dashed lines roughly separate out the SA mainland Tiger Snakes, from those on Kangaroo Island, and those on islands in the Nuyts Archipelago and Sir Joseph Banks Island group. The SA mainland samples generally experience higher annual mean temperatures (~15.7—17.3°C) compared to Tiger Snakes on the Nuyts Archipelago and Sir Joseph Banks islands (~13.5—16.3°C), which appears to be related to their genetic/epigenetic profiles (*Hpa*II PCoA1 values). The Pearson correlation coefficient (r = 0.652) shows a strong positive linear correlation.

**Figure 17.**
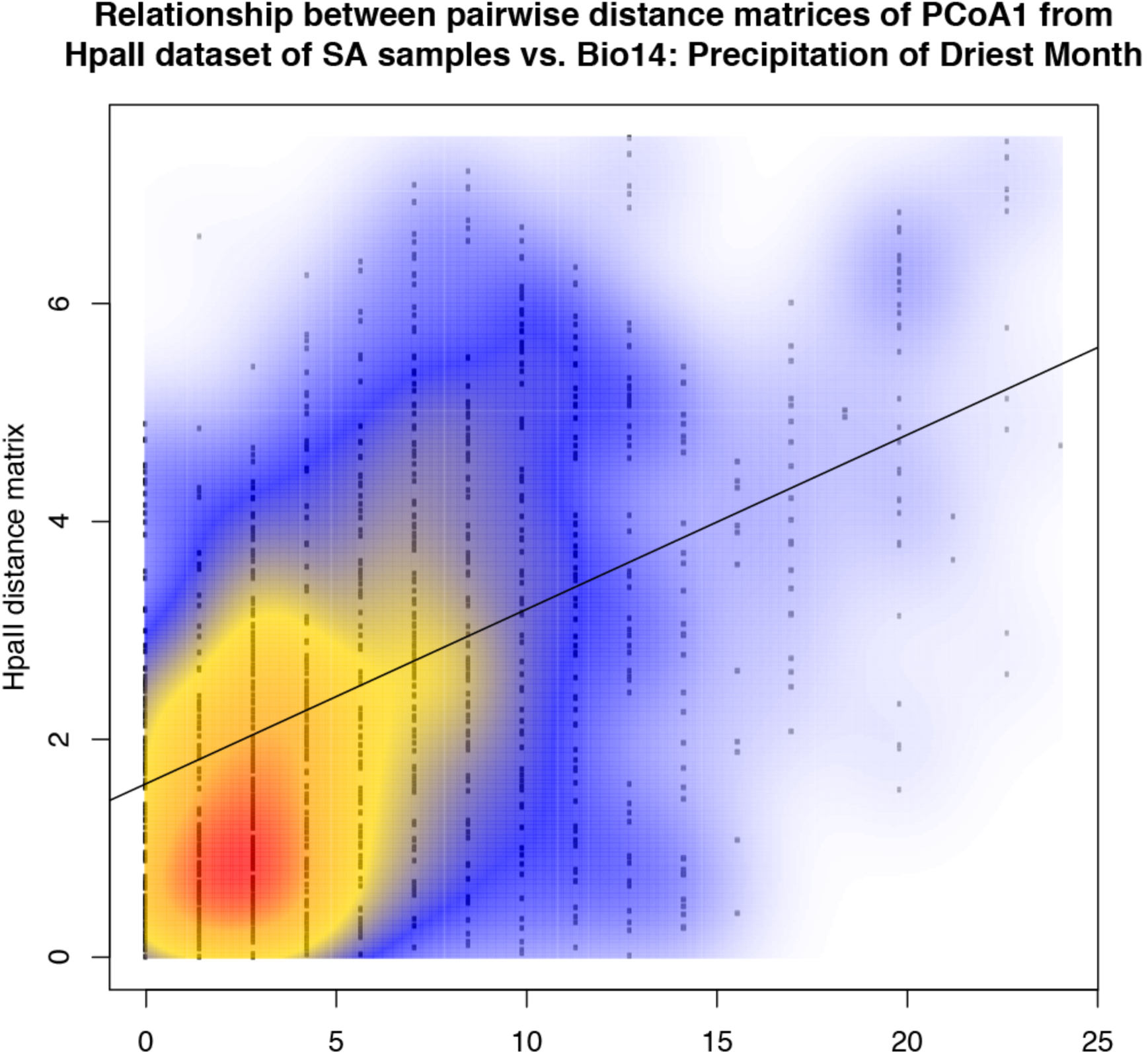
Mantel test showing correlation between pairwise distance matrix of *Hpa*II PCoA1 values and pairwise distance matrix of Bio14 variables (precipitation of the driest month) for Tiger Snakes from South Australian locations. The correlation is significantly different from random, with a p-value of 0.001 (Bonferroni corrected p-value of 5% significance level is 0.0026).

**Figure 18.**
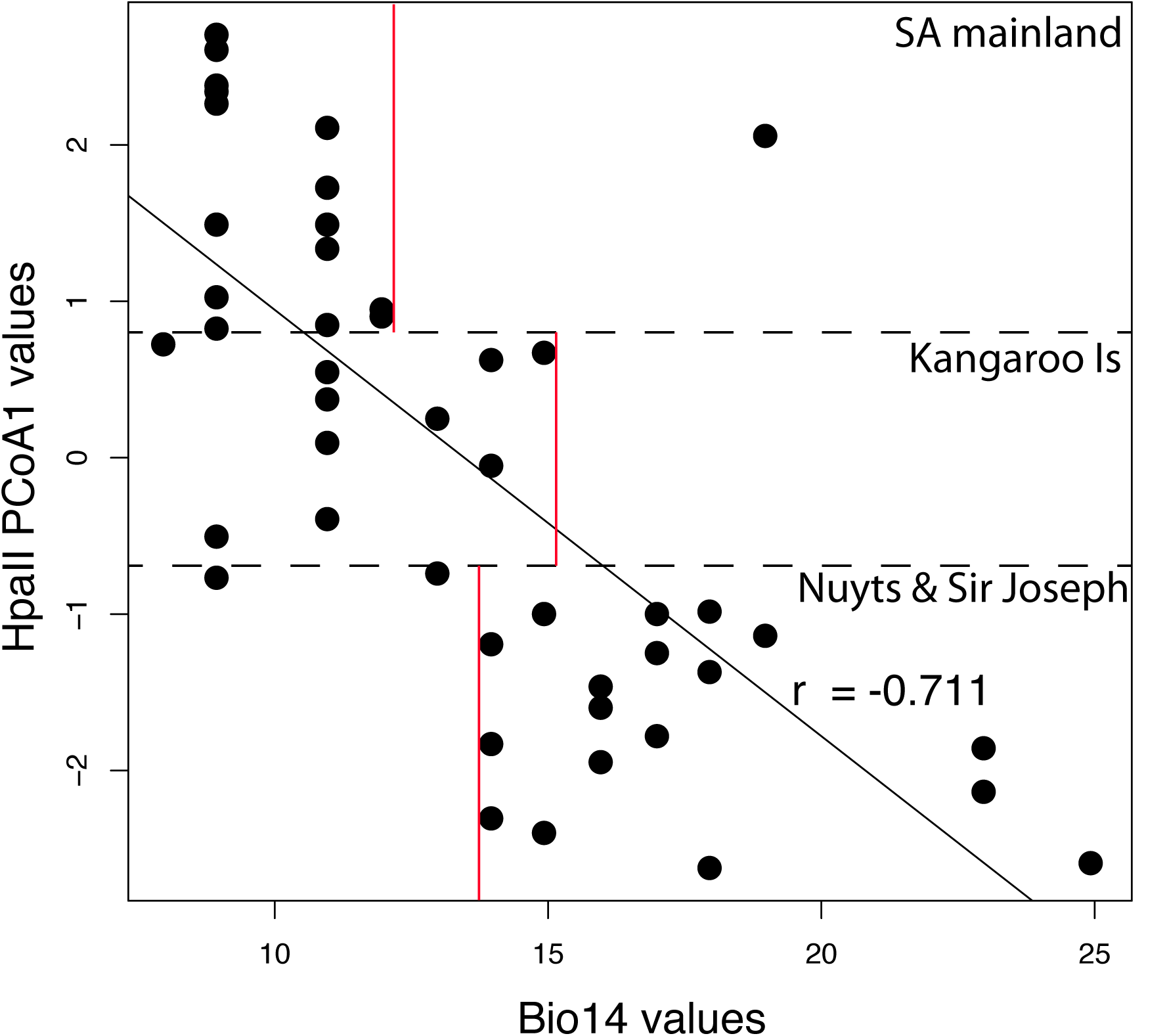
Scatterplot of raw *Hpa*II PCoA1 values vs. Bio14 (precipitation of the driest month) values for Tiger Snakes from South Australian (SA) locations. The horizontal dashed lines roughly separate out the SA mainland Tiger Snakes, from those on Kangaroo Island, and those on islands in the Nuyts Archipelago and Sir Joseph Banks Island group. The SA mainland samples generally experience lower precipitation levels in the driest month (< ~12mm) compared to Tiger Snakes on the Nuyts Archipelago and Sir Joseph Banks islands (> ~12mm), which appears to be related to their genetic/epigenetic profiles (*Hpa*II PCoA1 values). The Pearson correlation coefficient (r = -0.711) shows a strong negative linear correlation.

**Table 8.** Correlation between genetic/epigenetic distance *(Hpa*II PCoA1 pairwise distance matrix) and bioclimatic variables (pairwise distance matrix) for each SA sample.

| Variable | Mantel test p-value | Pearson correlation coefficient | Mainland values | Nuyts Archipelago & Sir Joseph Banks Island values |
| --- | --- | --- | --- | --- |
| **Bio1: Annual mean temp (°C)** | **0.001\*\*** | **0.652** | **> 15.8** | **< 16.3** |
| Bio2: Mean Diurnal Range | 0.779 | 0.083 |  |  |
| Bio3: Isothermality (BIO2/BIO7) (* 100) | 0.115 | 0.261 |  |  |
| Bio4: Temperature Seasonality | 0.574 | -0.015 |  |  |
| Bio5: Max Temperature of Warmest Month | 0.578 | 0.174 |  |  |
| Bio6: Min Temperature of Coldest Month | 0.068 | 0.215 |  |  |
| Bio7: Temperature Annual Range (BIO5-BIO6) | 0.737 | 0.018 |  |  |
| Bio8: Mean Temperature of Wettest Quarter | 0.001** | 0.458 | > 12.0 | < 13.5 |
| Bio9: Mean Temperature of Driest Quarter | 0.002* | 0.469 |  |  |
| Bio10: Mean Temperature of Warmest Quarter | 0.002* | 0.478 |  |  |
| Bio11: Mean Temperature of Coldest Quarter | 0.001** | 0.598 | > 12.0 | < 12.2 |
| Bio12: Annual Precipitation (mm) | 0.001** | -0.506 | < 540 | 280—800 |
| Bio13: Precipitation of Wettest Month | 0.008 | -0.371 |  |  |
| **Bio14: Precipitation of Driest Month** | **0.001\*\*** | **-0.711** | **< 12** | **> 14** |
| Bio15: Precipitation Seasonality | 0.111 | 0.186 |  |  |
| Bio16: Precipitation of Wettest Quarter | 0.008 | -0.405 |  |  |
| Bio17: Precipitation of Driest Quarter (mm) | 0.001** | -0.691 | < 46 | > 47 |
| Bio18: Precipitation of Warmest Quarter | 0.001** | -0.677 | < 47 | > 48 |
| Bio19: Precipitation of Coldest Quarter | 0.001** | -0.411 | 125—240 | 80—330 |

One asterisk (\*) signifies Bonferroni corrected p-values less than 5% significance level (0.0026), with two asterisks (\*\*) representing Bonferroni corrected p-values less than 2.5% significance level (0.0013).

## Discussion

The MSAP technique only detects methylation at 5’-CCGG-3’ sites and cannot discriminate between methylation and fragment absence when both cytosines are hypermethylated: therefore the level of genomic DNA methylation may be underestimated. The *Hpa*II and *Msp*I datasets also have the added limitation of possibly still retaining some genetic signal, which complicates the epigenetic patterns. Bearing in mind these inherent limitations of the technique, our study has explored the level and pattern of genome-wide 5’-CCGG-methylation in Tiger Snake island and mainland populations.

When we compare the purely nuclear fragment length (AFLP) data, we see similar patterns to that observed in mtDNA (Keogh et al. 2005): kangaroo island Tiger Snakes fall with SA mainland samples and sister to some of the island samples (Sir Joseph Banks Island group). The epigenetic (*Hpa*II) data adds an extra dimension to the information about the Kangaroo Island Tiger Snakes, in that they look like mainland populations both genetically (AFLP) and epigenetically (*Hpa*II). This could be because of the relatively large size of the island and/or the complex climate/habitat/prey types etc. The low levels of within-population epigenetic diversity on KI suggests that even given the more complex island environment, the Tiger Snakes on KI have all responded epigenetically in a similar way to each other and to those on the mainland. A large complex environment may allow enough heterogeneous habitats that behavioural adaptation *(i.e.* via migration to new environments) is favoured over epigenetic *in-situ* adaptation that is the only option in range and environmentally restricted habitats, such as those that occur on small isolated islands.

There is more epigenetic differentiation within the mainland population as a whole compared to that within the islands as a group again possibly because the environment on the mainland is more heterogeneous than across the islands (Table 5). Another suggestion that the epigenetic signal is related to complex environmental differences is the fact that the Nuyts Archipelago and Sir Joseph Banks Island group are similar genetically to each other but significantly different in epigenetic signal even though they have been isolated for similar amounts of time and from presumably the same source population (Table 4, 6 & 7).

Even taking into account some retained genetic signal in PC1 of the *Hpa*II PCoA plots, these plots show interesting patterns: 1) the SA mainland populations are quite distinct from the Nuyts Archipelago and Sir Joseph Banks islands in PC space, which is not seen in the purely genetic data; 2) the intra-population variation *(i.e.* degree of clustering of each island population) is smaller in the epigenetic PCoA than in the genetic PCoA; and 3) the correlation between epigenetic and geographic distance is stronger than between genetic and geographic distance. If there is retained genetic signal within PC1 of the *HpaII* PCoA (and of a similar nature to the genetic signal in the AFLP PCoA), then this retained genetic signal would in essence be adding noise to the epigenetic signal, suggesting the purely epigenetic signal should exhibit stronger separation between the mainland and island populations, even less intra-population variation in the island populations and stronger correlation between epigenetic distance and geographic distance than that observed. Unless the *Hpa*II cut sites present in this data occur purely by coincidence in adaptive genes, then these patterns suggest that the island populations have adapted through epigenetic mechanisms to their specific environments.

This idea of Tiger Snake adaptation to specific environments epigenetically is further reinforced by the very strong correlation between PC1 in the *Hpa*II PCoA with various temperature and precipitation bioclimatic variables. The size of each island, as well as the habitat and climate, all possibly play a role in how adaptable the Tiger Snakes need to be in terms of life history traits. There are distinct and quite different ecological niches on the islands of the Nuyts Archipelago and the Sir Joseph Banks Island group (colder and with more rain) compared to the mainland (warmer and with less rain). The islands of the Sir Joseph Banks Island group and Nuyts Archipelago are also generally small in size, with low profiles and little natural water (Robinson *et al.* 1996), suggesting that uncharacteristically hot and dry seasons might have a larger impact on the habitat and prey items of the Tiger Snakes on these islands. Furthermore, on small islands, Tiger Snakes may need to retain flexibility in body/head size development strategies in order to adapt to rapid and uncommon weather events, as well as in changes in habitat and/or prey as they can’t simply migrate to follow their preferred niche and prey items. For example, if seasonal storms wipe out mutton-bird nests, a lack of large high calorie prey items at a critical time of year may result in only small reptiles being available for neonates as they develop in their first year of age, with reduced gigantism one possible outcome through epigenetic down-regulation of growth genes.

Interestingly, a strong correlation occurs between epigenetic distance (PhiPT in *Hpa*II) and isolation age. This is counter-intuitive if epigenetic regulation of traits develops over short time frames and then these traits get genetically assimilation over longer time frames. Another possible explanation for this correlation between epigenetic distance and isolation age is that the retained genetic signal has been subject to neutral genetic drift, which might also appear as increasing epigenetic differentiation relative to isolation time. The fact that Tiger Snake epigenetic patterns are not that similar to the purely genetic patterns makes neutral drift an unlikely explanation however. Neutral drift has been posited as one explanation for significant correlation between genetic and epigenetic patterns in another MSAP study of wild animal populations, however they suggest that methylation variation being dependent on DNA sequence differences is a more likely explanation (Liu 2012). In the case of the Tiger Snake, the environmental variability found on the SA islands may be too extreme to select for a new optimal set of traits after isolation. This may necessitate continued epigenetic regulation of phenotypic plasticity over thousands of years rather than only an initial adaptive period after isolation followed by genetic assimilation as per the model explaining epigenetic regulation of genes as a method of rapid adaptation to novel environments.

## Conclusion

In conclusion, by randomly subsampling the genome and epigenome of the Tiger Snake we found that Kangaroo Island Tiger Snakes resemble those on the mainland both genetically and epigenetically to a large degree, possibly because both KI and the mainland are environmentally complex. More importantly, we have also shown that the SA island populations have adapted to their environment in terms of both specific temperature and precipitation levels. Future work in this sphere needs to use finer-scale methods to examine both the genetic and epigenetic signal across populations of the Tiger Snake. Furthermore, a closer examination of epigenetic regulation of specific genes involved in growth *(i.e.* for head and body size) and lipid metabolism are needed to see epigenetic differences related to prey item variance on the different islands and in comparison to the mainland Tiger Snakes.

